# Understanding p300-transcription factor interactions using sequence variation and hybridization

**DOI:** 10.1101/2021.12.16.472944

**Authors:** Fruzsina Hobor, Zsofia Hegedus, Amaurys Avila Ibarra, Vencel L. Petrovicz, Gail J. Bartlett, Richard B. Sessions, Andrew J. Wilson, Thomas A. Edwards

**Affiliations:** Astbury Centre for Structural Molecular Biology, University of Leeds, Woodhouse Lane, Leeds LS2 9JT, UK; School of Molecular and Cellular Biology, University of Leeds, Woodhouse Lane, Leeds LS2 9JT, UK; Department of Medical Chemistry, University of Szeged, Dóm tér 8, H-6720 Szeged, Hungary; School of Biochemistry, University of Bristol, Medical Sciences Building, University Walk, Bristol BS8 1TD, UK; BrisSynBio, University of Bristol, Life Sciences Building, Tyndall Avenue, Bristol BS8 1TQ, UK; School of Chemistry, University of Bristol, Cantock’s Close, Bristol BS8 1TS, UK; School of Chemistry, University of Leeds, Woodhouse Lane, Leeds LS2 9JT, UK

## Abstract

The hypoxic response is central to cell function and plays a significant role in the growth and survival of solid tumours. HIF-1 regulates the hypoxic response by activating over 100 genes responsible for adaptation to hypoxia, making it a potential target for anticancer drug discovery. Although there is significant structural and mechanistic understanding of the interaction between HIF-1α and p300 alongside negative regulators of HIF-1α such as CITED2, there remains a need to further understand the sequence determinants of binding. In this work we use a combination of protein expression, chemical synthesis, fluorescence anisotropy and isothermal titration calorimetry for HIF-1α sequence variants and a HIF-1α- CITED hybrid sequence which we term CITIF. We show the HIF-1α sequence is highly tolerant to sequence variation through reduced enthalpic and less unfavourable entropic contributions, These data imply backbone as opposed to side chain interactions and ligand folding control the binding interaction and that sequence variations are tolerated as a result of adopting a more disordered bound interaction or “fuzzy” complex.

## Introduction

The hypoxic response is crucial to cell survival; it needs to both rapidly adapt to subtle variations in, and fluctuating, oxygen levels, and, allow recovery from hypoxia.^1-3^ As low oxygen level is a universal hallmark of solid tumours, the ability to adapt to hypoxia is essential for their growth and survival.^4^ The hypoxic response is mediated by transcriptional activation of genes that facilitate either short term (e.g. increased vascular permeability, glucose transport) or long term adaptive mechanisms (such as angiogenesis);^5-7^ these processes are largely mediated by the transcription factor Hypoxia Inducible factor (HIF) 1.^5-7^ HIF-1 is responsible for the activation of over 100 genes that play essential roles in the hypoxic response and thus plays a role in tumour growth and survival, making it a potential target for anticancer drug discovery.^8-12^ Indeed, a number of approaches to target protein-protein interactions of HIF-1 have been explored.^11, 13-24^ HIF-1 is a heterodimer, consisting of two subunits, the constitutively expressed HIF-1β and the oxygen sensitive HIF-1α.^3^ Under normoxic conditions, HIF-1α undergoes hydroxylation leading to interaction with the E3 Ligase pVHL and degradation, whereas under hypoxic conditions this is suppressed resulting in accumulation and translocation of HIF-1α to the nucleus where it forms a heterodimer with HIF-1β and recruits transcriptional co-activators, such as p300.^8, 25-30^ The multidomain protein p300 and its paralogue CREB binding protein (CBP) are very similar in structure; they comprise a number of domains including the nuclear interaction domain (Nu), the CREB and MYB interaction domain (KIX), cysteine/histidine regions (CH/TAZ), a histone acetyltransferase domain (HAT) and a bromodomain (Br).^31-32^ The CH1 domain (which differs by only a few amino acids between p300 and CREB^33-34^ interacts with the C-TAD of HIF-1α. The CH1 domain has been shown to interact with a number of transcription factors including HIF-1α,^28, 33^ CREB-binding protein/p300-interacting transactivator with ED-rich tail (CITED 2),^35-36^ p53,^37^ NF-kB p65 subunit RelA,^38^ and, signal transducer and activator of transcription 2 (STAT2)^39^ through a range of recognition modes.^40^ Of particular interest, CITED2, is a negative feedback regulator that reduces HIF-1 transcriptional activity by competing for p300/CBP.^41-45^ HIF-1α and CITED2 have been reported to operate *via* a hypersensitive regulatory switch that exploits the properties of intrinsic disorder, similar p300/CBP binding affinities and a common LP(Q/E)L sequence mechanistically essential for binding, flanked by helical regions. CITED2 has been reported to displace HIF-1α from the surface of p300/CBP *via* transient ternary complex formation with both p300/CBP and HIF-1α followed by a subsequent shift in conformation resulting in a kinetic lock and prevention of the reverse process (i.e. displacement of CITED2 by HIF-1α).^46-47^ This provides a rationale as to why HIF- 1α transcriptional activity is sensitive to moderate CITED concentrations^41^ allowing effective negative feedback.

HIF-1α interacts with p300/CBP via its carboxy terminal transactivation domain (C-TAD). The solution structure of HIF-1α C-TAD in complex with p300/CBP was previously determined by NMR.^28, 33^ The CH1 domain of p300/CBP forms a rigid globular structure consisting of four α-helices (referred to here as α_1-4_), stabilised and constrained by three Zn atoms. The isolated C-TAD domain of HIF-1α is disordered in the absence of its binding partner. When bound to p300/CBP the HIF-1α C-TAD consists of three distinct α-helical regions and wraps around the p300/CBP CH1 domain^28^ (Fig. 1c-d, note in structure PDB ID: 1L3E^33^ the N-terminal region does not adopt a helical conformation). Several studies provide contradictory conclusions as to the importance of various regions and residues on HIF-1α C-TAD for p300/CBP.^17, 19, 48-49^ Mutational studies proposed key binding residues of HIF-1α;^48^ the *N-*terminal helix (HIF-1α_782-790_, also referred to as HIF-1α α_A_) has been shown to be less important for p300/CBP binding whilst the central and C-terminal helices (HIF-1α_797-805_ and HIF-1α_815-826_, also referred to as HIF-1α α_B and_ HIF-1α α_C_ respectively) of the HIF-1α C-TAD have been shown to be required for p300 recognition.^50^ HIF-1α_797-805_ bears two residues, Cys800 and Asn803, which can undergo post-translational modifications that modulate binding,^15, 49, 51-52^ and HIF-1α_815-826_ helix residues Leu818, Leu822 and Val825 are also considered important for binding.^48^ Additional HIF-1α_815-826_ helix residues that have been suggested to be important for recognition, include Asp823 and Gln824.^17, 19^ The potency of sequences derived from HIF-1α C-TAD (HIF-1α_776–826_, HIF-1α_786–826_ HIF-1α_788–822_ HIF-1α_776–813_) binding to p300/CBP were compared using fluorescence polarization.^49^ From this experiment it was concluded that the C-terminus of HIF-1α C-TAD is important for binding, in agreement with the mutagenesis studies.^33, 48^ Moreover, p300 sequence variants within the region that binds HIF-1α_815-826_ highlight its importance: whilst His349Ala and Leu376Met p300 variants showed minimal difference in HIF-1α affinity, a significant drop in potency was observed for the Ile400Met p300 variant;^50^ all these variants are found within the HIF-1α_815-826_ binding region with Ile400 closest to HIF-1α_815-826_. Site-directed mutagenesis in combination with kinetics measurements have been used to study the transition state for binding p300/CBP and the HIF-1α C-TAD: 17 HIF-1α C-TAD sequence variants were generated and binding assessed. Φ-Value binding analysis suggested that native hydrophobic binding interactions do not form at the transition state.^53^ HIF-1α Asn-803 hydroxylation was also shown to have a minimal destabilization effect. These data suggest the rate-limiting transition state is “disordered-like”, with subsequent co-operative formation of native binding contacts and replicates results observed for other p300/CBP CH1 interactions.^54^

**Figure 1.**
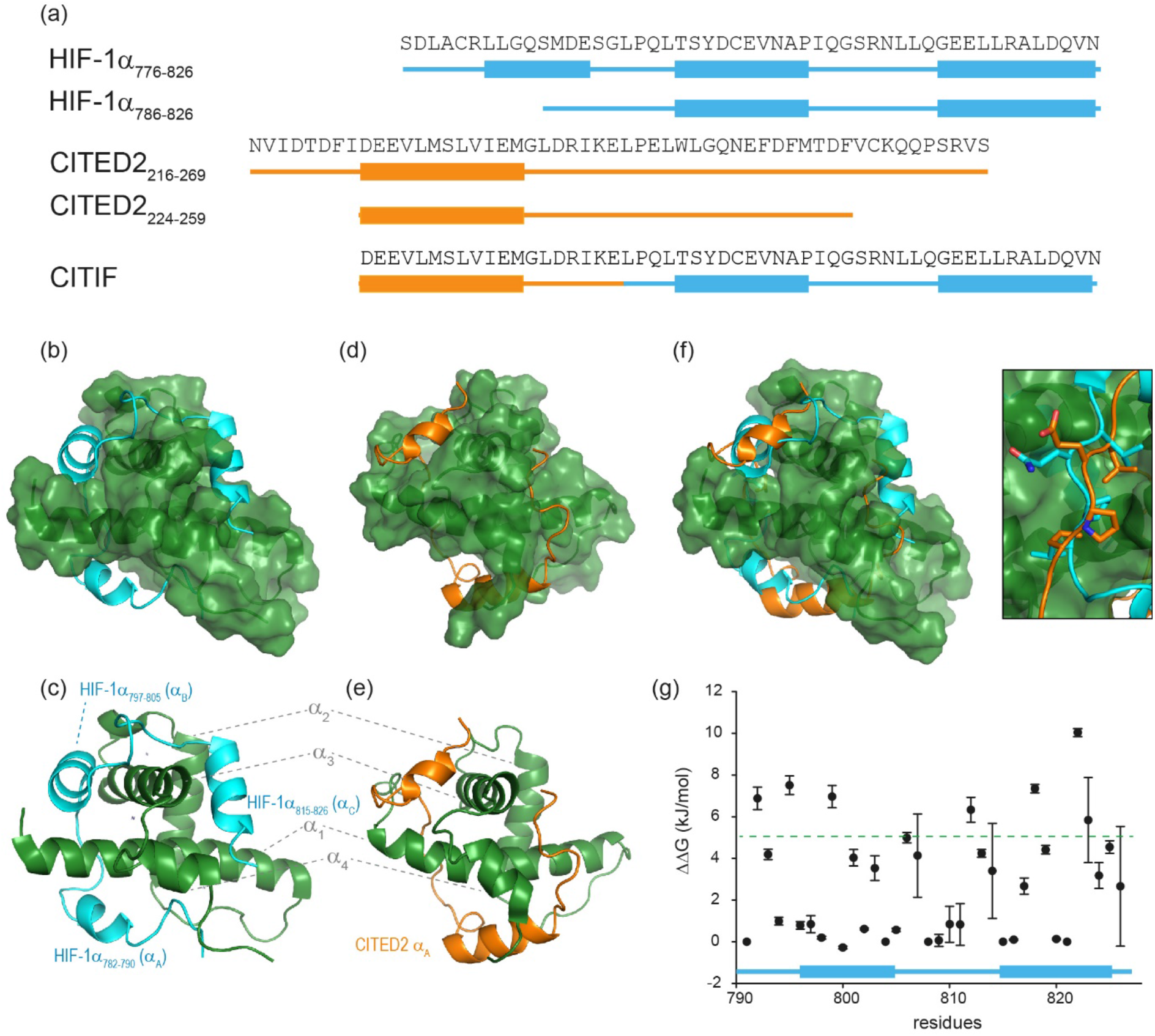
Sequences and structures of the p300 transcription factor complexes investigated in this work and binding free energy predictions on sequence determinants. (a) Sequence variants of HIF-1α and CITED2, helical regions are indicated by rectangles under the sequences. (b) Lowest energy structure from an NMR derived ensemble of the HIF-1α CTAD (cyan fold) and CBP(p300) CH1 domain (green surface) interaction (PDB ID: 1L8C); (c) same structure with HIF-1α C-TAD (cyan fold) and CBP(p300) CH1 domain (green fold); key regions are annotated for both HIF-1α and p300 with corresponding nomenclature used in Appling *et al*.,^47^ for clarity; (d) lowest energy structure from an NMR derived ensemble of the CITED2 (orange fold) and CBP(p300) CH1 domain (green surface) interaction (PDB ID: 1P4Q); (e) same structure with CITED2 (orange fold) and CBP(p300) CH1 domain (green fold); key regions are annotated for both CITED2 and p300 with corresponding nomenclature used in Appling *et al*.,^47^ for clarity; (f) overlay of the HIF-1α C-TAD (cyan fold) and CITED2 (orange fold) interactions with CBP(p300) CH1 domain (inset highlights the region where the conserved LPE(Q)L residues interact); (g) results of hot residue prediction using in silico alanine scanning (BUDE, 20 lowest energy structures from the NMR ensemble used in the prediction, circles denote average predicted ΔΔG, error bars the standard deviation).

HIF-1α (residues 776–826) and CITED2 (residues 216–269) recognize partially overlapping binding sites on p300/CBP (Fig. 1d-f). The helices of HIF-1α and CITED2 and their conserved LP(Q/E)L motifs bind to the same surfaces of the p300/CBP CH1 domain. The region of CITED2 that is C-terminal to the LPEL motif binds in an extended conformation in the same site as the HIF-1α_797-805_ helix.^35-36^ Despite this significant structural and mechanistic understanding of transcription factor p300/CBP interactions, there is a need to further understand the determinants of binding at a sequence level. Motivated by our recent studies on the effects of the HIF-1α truncation on the HIF-1α/p300 interaction,^21, 50^ identification of peptide and non-antibody binding proteins through selection methods,^50^ and, development of designed HIF-1α/p300 inhibitors^18, 21, 23, 55^ we sought to understand those determinants. We used a combination of protein expression, chemical synthesis, fluorescence anisotropy and isothermal titration calorimetry to probe the binding of HIF-1α sequence variants, CITED2 and a HIF-1α-CITED2 hybrid sequence (which we term CITIF; Fig. 1a) to the p300 CH1 domain (residues 330-420, hereinafter referred as p300). Our results point to an interaction that is remarkably tolerant to sequence variation, despite a high degree of sequence conservation across species.^28^ The parent interaction is enthalpically very favourable and entropically unfavourable; it seems to tolerate sequence variation through reduced enthalpic and less unfavourable entropic contributions, features which support a hypothesis whereby interactions between ligand (HIF-1α) and protein (p300) exploit a combination of non-covalent contacts between the HIF-1α backbone (as opposed to side-chains) and the surface of well folded p300 CH1 domain, along with HIF-1α folding, driven by transient side-chain contacts and long range electrostatic interactions to derive binding free energy. Adopting a more disordered bound interaction or “fuzzy” complex is consistent with the observed changes in thermodynamic signature and might account for the broadly tolerated sequences.

## Results and Discussion

### HIF-1α single sequence variations have little effect on p300 binding affinity

We previously developed BUDE AlaScan as predictive tool to identify hot residues and experimentally validated it for α-helix and β-strand mediated interactions.^56-57^ In those cases the interaction was localized within a single helix or strand in at least one of the interacting partners. The extended nature of the HIF-1α/p300 interaction afforded an opportunity to test the capabilities of *in silico* alanine scanning where affinity may be dispersed across a larger number of amino acid residues (for comparison, the NOXA/MCL-1 interaction has MCL-1 binding affinity *K*_D_ ∼100nM with 19 residues in NOXA as opposed to HIF-1α with similar *K*_D_ but 42 residues). BUDE AlaScan can predict changes in ΔΔG of binding upon introducing single or multiple alanine variations in one of the interacting partners when compared to the binding energy of the wild-type protein; in this case for HIF-1α using the HIF-1α/p300 NMR derived ensemble (PDB ID: 1L8C). This analysis (Fig. 1g) predicted key determinants of the HIF-1α binding to be dispersed across the whole sequence with several residues in both HIF-1α_797-805_ and HIF-1α_815-826_ showing ΔΔG > 4.2 kJ/mol (the threshold for a hot residue).^57-59^ A number of these e.g. L792 and L822 show ΔΔG >> 4.2 kJ/mol ΔΔG with small standard deviation implying those positions are indeed important for p300 binding while other residues with smaller values and greater standard deviation, like D823 were less clear cut. ROBETTA^60^ provided similar data (see ESI, Fig. S1).

To experimentally compare the predictions, we carried out an *in vitro* biophysical study of several HIF-1α sequence variants. We assessed predicted hot residues and their interactions with p300 using the NMR structure to visualize the structural basis behind the predictions (See ESI, Fig. S2). These analyses helped refine a first series of alanine variants to prepare. We did not consider HIF-1α_782-790_ variants given prior studies which had established little overall effect from the presence/absence of these 8-10 residues.^50^ HIF-1α_776-826_ sequence variants were recombinantly prepared based on the predictions to test their binding to the recombinantly prepared p300 (Fig. 2). Given the length of the peptide (42 residues), this was considered advantageous as it obviates the need to chemically synthesise, label and purify multiple variants. As the C-TAD domain is unstructured in isolation it was recombinantly expressed as a fusion protein with GFP. The green-fluorescent protein (GFP) tag was used for fluorescence anisotropy experiments to determine the binding affinity of HIF-1α_776-826_ C-TAD variants to p300. As the CH1 domain of p300 is a small domain of 11 kDa it was recombinantly expressed as a fusion protein with GST to increase its size and thus the signal to noise in the FA experiments. We established an assay where the interaction between GFP-tagged wt HIF-1α_776-826_ and GST-tagged p300 was monitored by FA, using the fluorescence of GFP. As GST dimerises, we assume the stoichiometry of the interaction is 2:2. (Fig. 2a). To demonstrate that binding between HIF-1α and p300 was not affected by GST a control experiment was performed using GFP-HIF-1α_776-826_ and p300 with the GST tag cleaved; although the change in anisotropy signal was lower (consistent with the lower mass of the complex in the absence of GST) the determined *K*_D_ was comparable between the two experiments (Fig. S3). Similarly, ITC experiments for the binding of GFP-HIF-1α_776-826_ and HIF-1α_776-826_ to p300 were comparable (see later).

**Figure 2.**
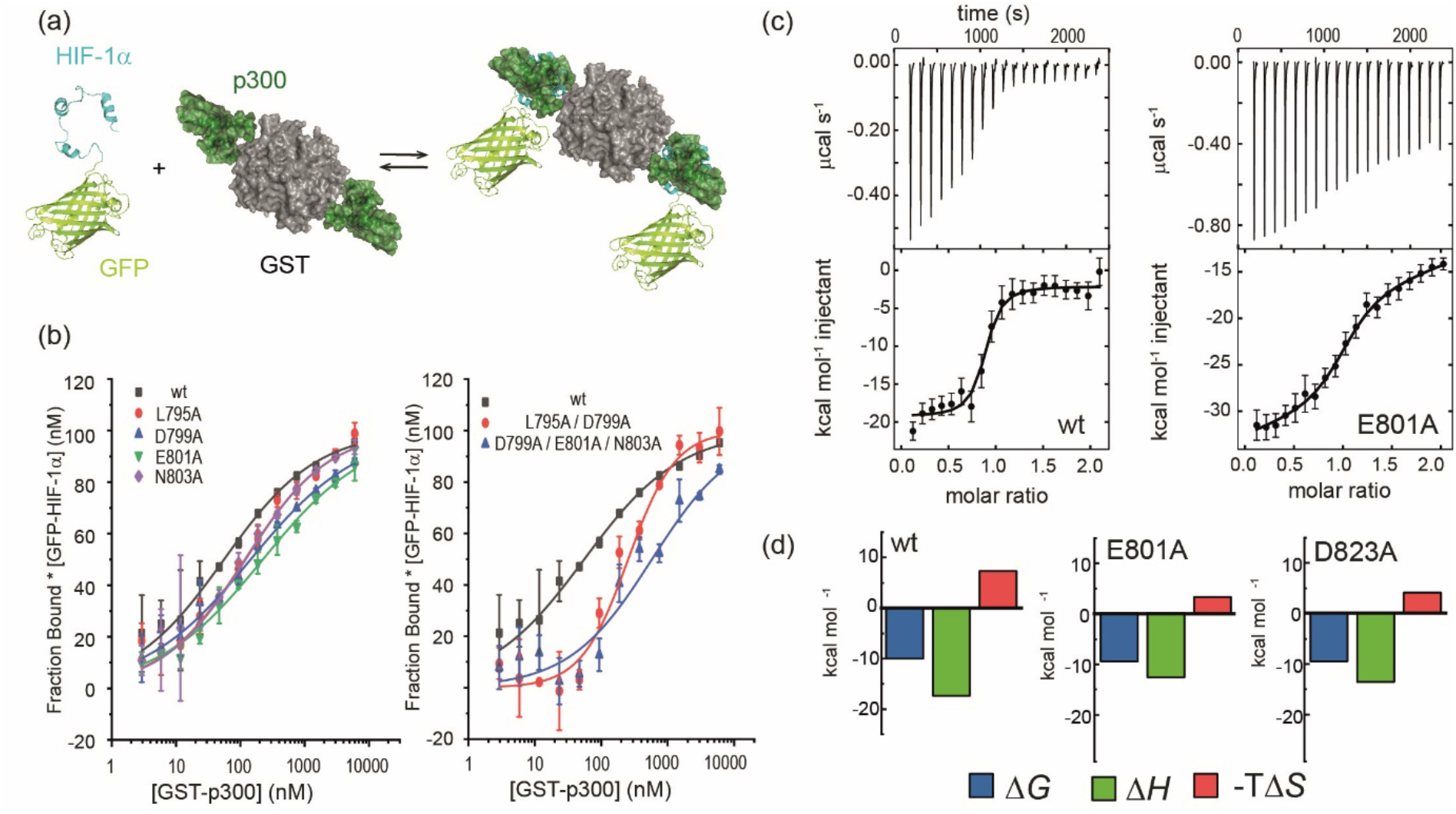
Biophysical analyses on the effects of HIF-1α sequence variant p300 binding affinity. (a) schematic depicting the equilibrium for interaction of GFP-HIF-1α_776-826_ variants and GST-p300 as studied by fluorescence anisotropy; (b) representative fluorescence anisotropy titration data for sAV and mAV HIF-1α_776-826_ peptides interacting with p300 (25 mM Tris-HCl, 150 mM NaCl, 1mM DTT, pH 7.4); (c) raw ITC data (upper) and fitted thermogram (lower) for the interaction of HIF-1α_776-826_ peptides with p300 (37°C in 25 mM Tris-HCl, 150 mM NaCl, 1mM DTT, pH 7.4) using 10 µM p300 in the cell and 100 µM HIF-1α_776-826_ variant in the syringe; (d) thermodynamic signatures for each interaction.

After establishing the assay, selected single alanine HIF-1α CTAD variants (sAVs) were prepared and their binding affinity to p300 was tested. Results for these experiments are shown in Figure 2b, Table 1 and Fig. S4. Our data show there is a limited impact upon the binding to p300 for single alanine variations introduced into HIF-1α_797-805_, HIF-1α_815-826_ or the LPQL sequence that shares homology with CITED2 (≤ 4 fold maximal difference). This contrasts with the work of Lindström who identified L792A, L795A and L818A as hot residues (alongside L812A and L813A, which were not considered here), although these were derived from Φ-value binding analysis using tryptophan fluorescence and may reflect transition state effects upon binding.

**Table 1.**
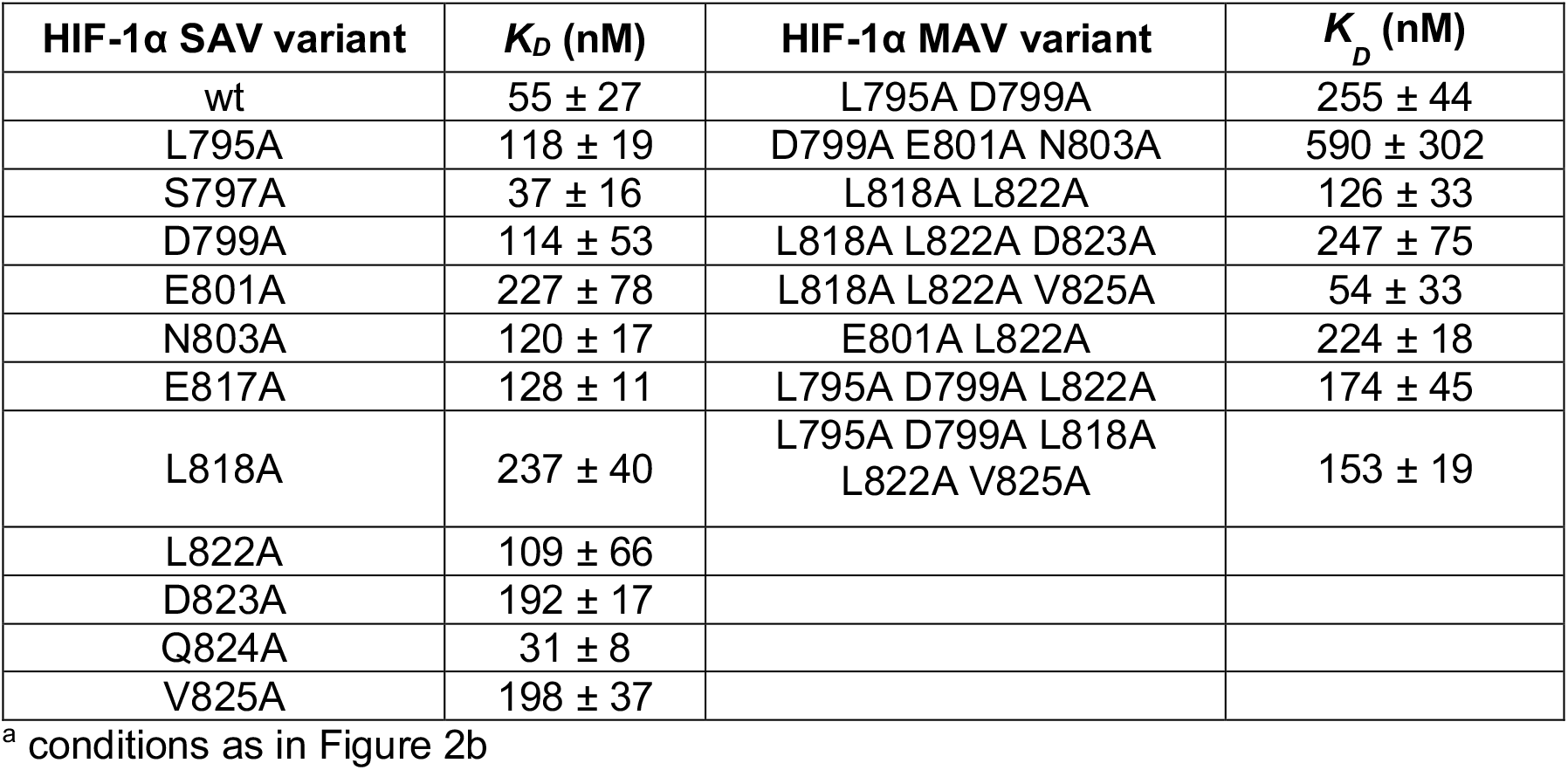
Dissociation constants for HIF-1α_776-826_ C-TAD single alanine mutant variants binding to p300

| HIF-1α SAV variant | $K_D$ (nM) | HIF-1α MAV variant | $K_D$ (nM) |
| --- | --- | --- | --- |
| wt | 55 ± 27 | L795A D799A | 255 ± 44 |
| L795A | 118 ± 19 | D799A E801A N803A | 590 ± 302 |
| S797A | 37 ± 16 | L818A L822A | 126 ± 33 |
| D799A | 114 ± 53 | L818A L822A D823A | 247 ± 75 |
| E801A | 227 ± 78 | L818A L822A V825A | 54 ± 33 |
| N803A | 120 ± 17 | E801A L822A | 224 ± 18 |
| E817A | 128 ± 11 | L795A D799A L822A | 174 ± 45 |
| L818A | 237 ± 40 | L795A D799A L818A<br>L822A V825A | 153 ± 19 |
| L822A | 109 ± 66 | | |
| D823A | 192 ± 17 | | |
| Q824A | 31 ± 8 | | |
| V825A | 198 ± 37 | | |

<sup>a</sup> conditions as in Figure 2b

We carried out ITC measurements for several variants to verify the results of the fluorescence anisotropy measurements (Fig. 2c and ESI, Fig. S5). The interaction of HIF-1α_776-826_ with p300 is characterized by a large favourable enthalpy of interaction and opposing unfavourable entropy of interaction. The dissociation constant was similar for all the tested variants and the thermodynamic signature shifted toward less favourable enthalpic contributions compensated by more favourable entropy (Figure 2c-d, Table 2.) The removal of a transient charge reinforced interaction (E801A and D823A) may increase the local flexibility of the structure resulting in the observed, less unfavourable entropy. This implies that HIF-1α can adjust its interaction with p300 to achieve optimal affinity also for the variants, which emphasizes the requirement to occupy the surface through ‘fuzzy’ interactions rather than specific contacts.

**Table 2.** Thermodynamic parameters for the binding of GFP-HIF-1α_776-826_ variants to p300. 68% confidence intervals are shown in brackets (conditions as in Figure 2c)

| <sup>a</sup> | $K_D$ nM | $\Delta H$<br>kcal mol<sup>-1</sup> | $\Delta S$<br>cal mol<sup>-1</sup> K<sup>-1</sup> |
| --- | --- | --- | --- |
| HIF-1α | 86<br>(32 – 182) | -17.3<br>(-18.1 to -16.6) | -23.6 |
| HIF-1α E801A | 359<br>(284 - 444) | -12.2<br>(-12.6 to -11.9) | -9.5 |
| HIF-1α D823A | 369<br>(264 – 496) | -13.0<br>(-12.5 to -13.6) | -12.2 |

<sup>a</sup> we also prepared an L792A variant which exhibited nM affinity although the quality of data fitting impedes a more detailed analyses of thermodynamic parameters in this instance (see Figure S5).

**Table 3.** Thermodynamic parameters for the binding of HIF-1α, CITED2 and CITIF peptides to p300

|  | Synthetic Peptides<sup>a</sup> |  |  | Expressed Peptides<sup>b</sup> |  |  |
| --- | --- | --- | --- | --- | --- | --- |
| | $K_D$ (nM) | $\Delta H$ kcal mol<sup>-1</sup> | $\Delta S$ cal mol<sup>-1</sup> K<sup>-1</sup> | $K_D$ (nM) | $\Delta H$ kcal mol<sup>-1</sup> | $\Delta S$ cal mol<sup>-1</sup> K<sup>-1</sup> |
| HIF-1α<sub>786-826</sub> | 42.9<br>(38.3 - 47.6) | -25.8<br>(-25.6 to -26.1) | -50 | 88.3<br>(57.0 -132.4) | -32.7<br>(-31.5 to -33.9) | -73.2 |
| HIF-1α<sub>776-826</sub> | 52.2<br>(49.2 - 55.8) | -22.9<br>(-22.8 to -23.1) | -41 | 82.4<br>(55.0 -119.5) | -22.2<br>(-21.5 to -22.9) | -39.2 |
| CITED<sub>224-259</sub> | 9.1<br>(6.5 - 12.2) | -12.4<br>(-12.2 to -12.7) | -3.6 | 41.9<br>(28.9 - 59.5) | -12.4<br>(-12.0 to -12.7) | -6.2 |
| CITED<sub>216-269</sub> | 26.3<br>(20.8 - 33.1) | -13.2<br>(-12.9 to -13.4) | -10.0 | 18.8<br>(12.2 - 27.6) | -13.3<br>(-13 to -13.5) | -7.5 |
| CITIF | 11.3<br>(9.3 - 13.7) | -22.5<br>(-22.3 to -22.7) | -37.8 | 15.4<br>(10.4 - 21.8) | -18.3<br>(-17.9 to -18.7) | -23.3 |

<sup>a</sup> conditions as in Figure 4
<sup>b</sup> conditions as given in Figure S6

### HIF-1α multiple sequence variations do not affect p300 binding affinity

To assess the extent to which sequence variations could confer additive effects on binding affinity, different, structurally relevant combinations (i.e. with the highest combined predicted ΔΔG values) of alanine variations were introduced into the HIF-1α_776-826_ C-TAD and their binding affinity determined (Fig. 3, Table 1 and ESI Fig. S4). The experimental data for these multiple alanine variants (mAVs) clearly shows that variations of two or three predicted hot residues either in HIF-1α_797-805_ or HIF-1α_815-826_ are generally insufficient to abrogate p300 binding. Even introducing variations in two helices (e.g. E801A L822A) simultaneously did not increase the *K*_D_ significantly; variants generally maintained affinity to p300 although for some mAVs (e.g., D799A-E801A-N803A; L795A-D799A; L818A-L822A-D823A and E801A L822A), there appears to be some loss in potency. Lower net negative charge of the TADs influences the long-range electrostatic interactions leading to lower association rates,^61^ which, in part may explain the decreased binding affinity of some of these mAVs. Collectively, these data further support a conclusion that the HIF-1α/p300 interface is fuzzy in nature; the plasticity in the interaction allows for signficiant sequence variation in the HIF-1α C-TAD with loss of one side chain likely to be compensated for by interactions of other side chains, possibly augmented by interactions of the backbone with the p300 surface.

**Figure 3.**
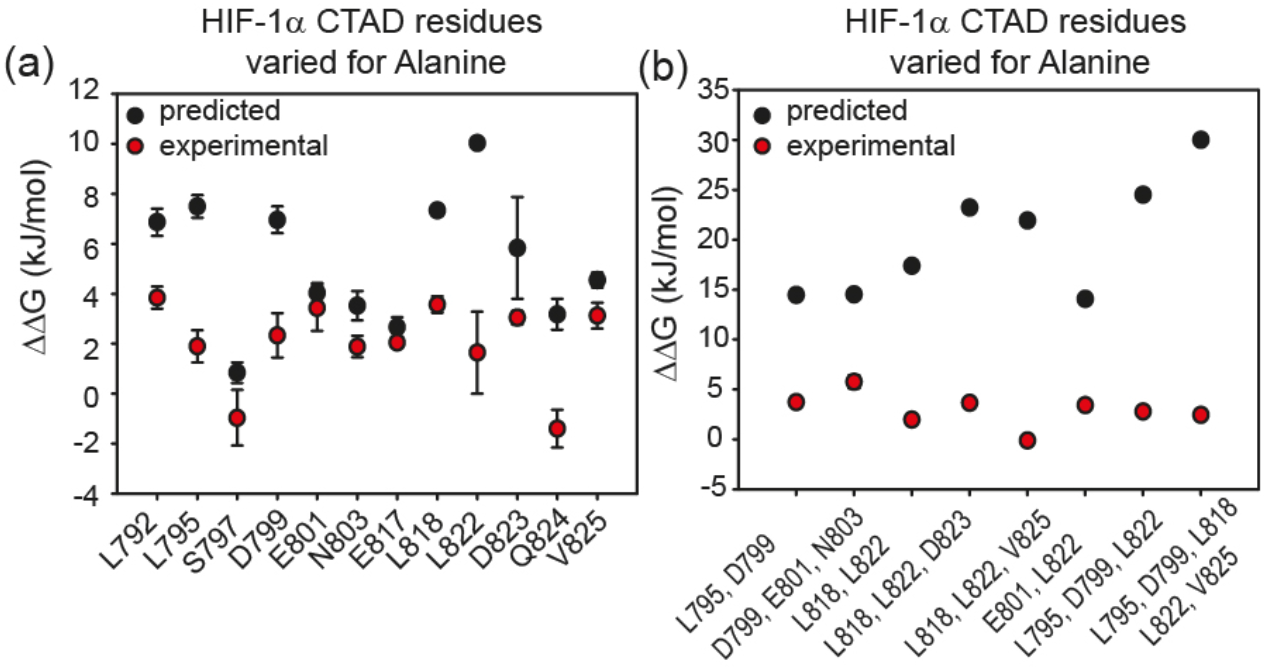
Comparison of predicted and experimental ΔΔG values for: (a) single alanine variant (sAV) and (b) multiple alanine variants of HIF-1α C-TAD for binding to p300. ΔΔG values were derived from FA measurements, except for L792A for which ITC data was used.

**Figure 4.**
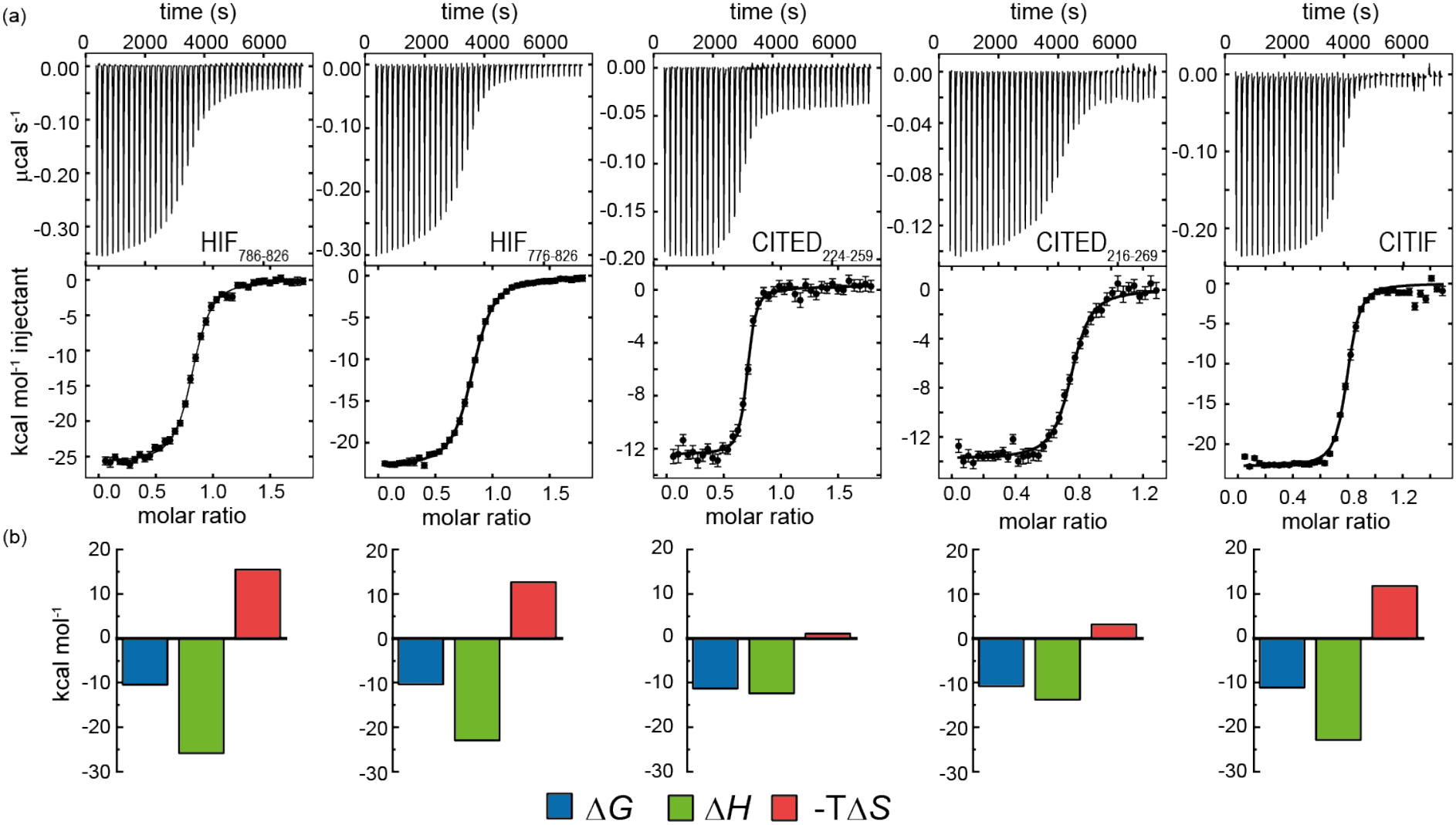
(a) Isothermal titration calorimetry data for the interaction of chemically synthesized HIF-1α, CITED2 and CITIF peptides with p300. Raw ITC (upper) data and fitted thermogram (lower) (40 mM sodium phosphate, pH 7.5 100 mM NaCl, 1 mM DTT buffer using 5 µM protein in the cell and 60 µM ligand in the syringe at 35°C); (b) Thermodynamic signatures for each interaction.

### Comparison of predicted and experimental variant HIF-1α/p300 binding affinities

The experimental determined values for sAVs and mAVs do not agree fully with the predictions (Fig 3). It should be noted that the predictions (both using BUDE and ROBETTA) did not identify particularly large sAV ΔΔG values > 8 kJ mol^-1^ and our earlier work highlighted the challenges in accurately predicting absolute values of ΔΔG using fast methods which are well suited to a yes/no indicator.^56-57^ Overall, the comparison between prediction and experiment for sAVs reveals the predictions overestimate the change in affinity, although there is still a moderate effect for most predicted hot-residues. Comparison of the prediction and experiment for mAVs reveals more pronounced differences; the additive combination of sequence variations is predicted to be significant (> 15 kJ/mol in many cases), yet minimal effects are observed for as many as five simultaneous sequence variations. This is consistent with the interaction becoming more fuzzy upon sequence variation to compensate for loss of side-chain interactions, a property not assessed in predictive alanine scanning.

Taken together our results suggest that interaction of some of the side-chains from each helix of HIF-1α are sufficient to maintain nanomolar affinity for p300; as the three helices wrap around p300; varying one or two positions is not sufficient to disrupt the binding, implying a high degree of chelate co-operativity (observed in our earlier truncation studies)^50^ and dispersal of binding energy across the sequence. As noted above for the thermodynamic analyses, the large favourable enthalpy and unfavourable entropy of binding for the native HIF-1α/p300 interaction together with the well tolerated sequence variation and observed enthalpy-entropy compensation for variants predicted to have diminished p300 affinity points to a key role of backbone or long range electrostatic interactions (which are not explored using computational alanine scanning) and transcription factor folding to generate binding energy. Such behaviour and any potential decrease of unfavourable steric contacts would accommodate sequence variation where the variant bound complex is more disordered relative to the native bound complex.

### CITED2 has higher affinity than HIF-1α for p300 and exhibits a sequence dependent competition mechanism

We hypothesized that it would be possible to enhance the affinity of HIF-1α for p300 by hybridising key regions of both CITED2 and HIF-1α (see later). We first measured the affinity of the parent peptides. A particular difficulty in comparing these peptide sequences is the different length used in different studies.^28, 33, 35-36^ We therefore considered HIF-1α_786-826_, HIF-1α_776-826_ CITED2_224-259_ and CITED2_216-269_ for these analyses and studied binding to p300 using ITC. Initially we expressed these as GFP fusion proteins and cleaved the tag, however the peptides all contained four residues from the PreScission protease sequence (ITC data given in the ESI Fig. S6).

Subsequently we also developed a chemical synthesis of the peptides bearing an N-terminal acetamide and C-terminal amide (see ESI). In general both sets of reagents gave similar data in terms of *K*_D_ - one notable exception is CITED_224-259_ which gave a *K*_D_ four-fold lower in magnitude for the expressed peptide relative to the chemically synthesized peptide. It may be that the four residues (Gly-Pro-Gly-Ser) remaining from the Prescission protease cleavage or free N-terminus interfere with p300 recognition. Support for this hypothesis is strengthened by the fact that both HIF-1α sequences also have weaker affinity (although not as pronounced) in comparison to the synthetic peptides. Overall, the CITED2 peptides have slightly higher p300 affinity than the HIF-1α peptides. This differs from observations reported by Berlow *et al*. who observed identical *K*_D_s of 10 nM for HIF-1α_776-826_, 10 nM CITED2_216-269_ both labelled with Alexafluor 488/595. In this prior work, a variety of biophysical and NMR methods were used to show that despite similar potencies, CITED2 effectively displaces HIF- 1α from the surface of p300 *via* transient ternary complex formation with both p300 and HIF-1α followed by a subsequent shift in conformation resulting in a kinetic lock and suppression of the reverse process (i.e. displacement of CITED2 by HIF-1α).^46^ Although the NMR experiments were performed at higher concentration, the fluorescent experiments used to determine affinity were performed at lower concentrations; the fluorescent labels and their positions may influence the equilibrium. The ITC data on unlabelled peptides which we report here suggest that the moderate preference for interaction of CITED2 with p300 over HIF-1α may incorporate a thermodynamic aspect and not exclusively derive from kinetic factors.

Competition experiments indicated that CITED2 and HIF-1α bind with negative cooperativity to p300 with a mechanism depending on the length of the peptide (Fig. 5). The apparent *K*_D_ for CITED_224-259_ displacing HIF_786-826_ from p300 is 665 nM (Δ*H* = 9.35 kcal/mol) and for HIF_786-826_ in the reverse process is 4.2 µM (Δ*H* = -5.6 kcal/mol), which is close to the expected values if the ligands bind competitively (Fig. S7, Table S1). Global analysis using a competitive binding model resulted in thermodynamic parameters that were concurrent with the direct titration experiments (Table 4.). On the other hand, CITED_216-269_ displaces HIF-1α_776-826_ more effectively with an apparent *K*_D_ of 43 nM (Δ*H* = 6.76 kcal/mol) which indicated cooperative binding. This differs significantly from the value reported by Berlow et al.; they report that the apparent *K*_D_ for CITED2 against the p300/HIF-1α = 0.2 nM, a 50-fold lower value than *K*_D_ determined for the direct CITED2/p300 interaction. Global analysis of the ITC data suggested a ternary complex formation with a ΔΔG of 0.31 kcal/mol and ΔΔH 20.5 kcal/mol (Table 4.) This data is consistent with the model where CITED2 and HIF-1α bind simultaneously to p300 forming a ternary complex which is destabilized by unfavourable enthalpy change and compensated by favourable entropic contributions. The sequence dependence of the competition mechanism suggests that CITED2_216-269_ contains the key residues that are responsible for the allosteric effect favouring unidirectional displacement of HIF-1α by CITED2. This is supported by studies on a C terminally truncated construct (CITED2_216-248_), which despite having lower affinity to p300 (*K*_D_ = 303 nM), displaces HIF-1α_776-826_ with similar efficiency to the higher affinity CITED2_224-259_ (*K*_d,app_ = 2 µM, Table S1-2). Furthermore, the removal of the N terminal eight residues (CITED2_224-248_) results in significantly decreased affinity, highlighting the importance of these residues for binding (Figure S8, Table S2).

**Figure 5.**
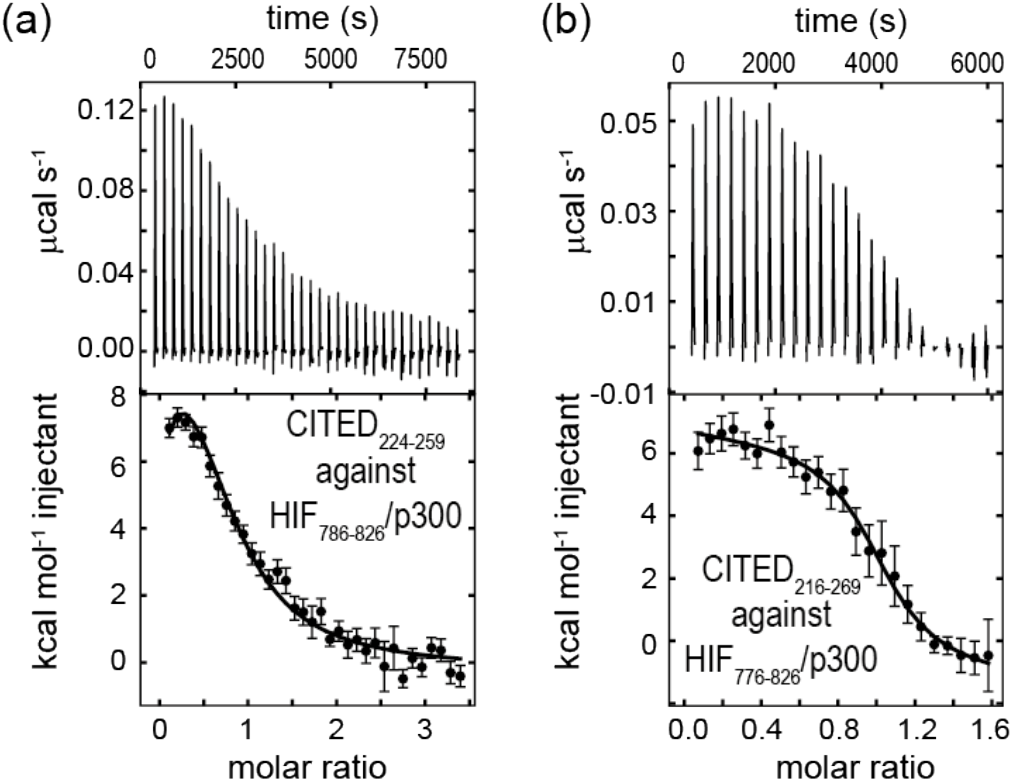
Competition ITC experiments (a) Raw ITC (upper) data and fitted thermogram (lower) for CITED2_224-259_ titrated against the p300/HIF-1α_786-826_ complex and (b) CITED2_216-269_ titrated against the p300/HIF-1α_776-826_ complex (40 mM sodium phosphate, pH 7.5 100 mM NaCl, 1 mM DTT buffer at 35°C). Complexes were prepared by titrating p300 with the competitor ligand until it reached saturation, which resulted in 1.2-1.6 equivalent ligand in the cell. Concentrations and exact molar ratios are listed in Table S2. The thermograms were fitted globally including datasets for direct titrations.

**Table 4.**
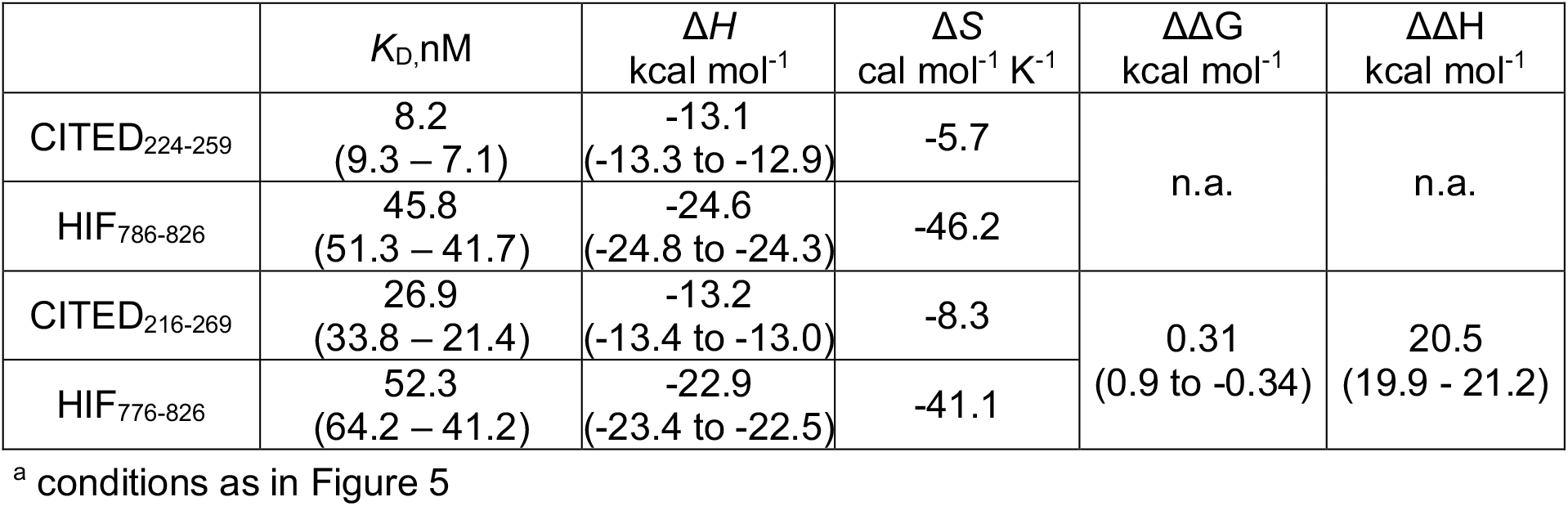
Thermodynamic parameters extracted from the global fitting of the competition titrations. The CITED2_224-259_ / HIF-1α_876-826_ system was fitted using a competitive binding model, the CITED2_216-269_ / HIF-1α_776-826_ system was fitted including a fit for ΔΔG and ΔΔH. 68% confidence intervals are shown in brackets.

| | $K_D$, nM | $\Delta H$<br>kcal mol<sup>-1</sup> | $\Delta S$<br>cal mol<sup>-1</sup> K<sup>-1</sup> | $\Delta\Delta G$<br>kcal mol<sup>-1</sup> | $\Delta\Delta H$<br>kcal mol<sup>-1</sup> |
| --- | --- | --- | --- | --- | --- |
| CITED<sub>224-259</sub> | 8.2<br>(9.3 – 7.1) | -13.1<br>(-13.3 to -12.9) | -5.7 | n.a. | n.a. |
| HIF<sub>786-826</sub> | 45.8<br>(51.3 – 41.7) | -24.6<br>(-24.8 to -24.3) | -46.2 |  |  |
| CITED<sub>216-269</sub> | 26.9<br>(33.8 – 21.4) | -13.2<br>(-13.4 to -13.0) | -8.3 | 0.31<br>(0.9 to -0.34) | 20.5<br>(19.9 - 21.2) |
| HIF<sub>776-826</sub> | 52.3<br>(64.2 – 41.2) | -22.9<br>(-23.4 to -22.5) | -41.1 |  |  |

<sup>a</sup> conditions as in Figure 5

### A HIF-1α-CITED2 hybrid – CITIF – has comparable p300 binding affinity to CITED2, but exhibits intermediate enthalpic and entropic signature to those of the parent HIF-1α and CITED2 sequences

A hybrid sequence (CITIF) was designed containing an N-terminal fragment of CITED2 (224-243) and a C-terminal fragment of HIF-1α (792-826) fragment. Expressed and chemically synthesized peptides were tested with both giving a *K*_D_ slightly higher than the HIF-1α sequences and comparable to the CITED2 sequences (Table 2). A fluorescence anisotropy competition assay established that this hybrid sequence competes with HIF-1α for binding to p300, supporting the hypothesis that CITIF reproduces key binding features of both HIF-1α and CITED2 (Fig. S9). Whilst both the HIF-1α sequences were shown to have strongly favourable p300 binding enthalpies and strongly unfavourable p300 binding entropies, in contrast, both CITED2 sequences were shown to have much less favourable p300 binding enthalpies, and much less unfavourable p300 binding entropies. The CITIF sequence exhibited p300 binding enthalpies and entropies intermediate between those observed for HIF-1α and CITED2. We obtained co-crystals of p300 in complex with CITIF and solved the structure at 2Å resolution (Fig. 6, Table S3). The structure shows that residues corresponding to CITED2 and HIF-1α bind simultaneously, occupying their native binding sites and reproducing most of the native contacts with the protein (Fig S11.), in line with the thermodynamic signature we observed for CITIF binding. Similarly to the CITED2-HIF-1α fusion peptide/CBP complex (PDB: 7LVS, Fig S12) recently reported by Appling *et al*., the N-terminal p300_345-373_ helix (α_1_) is straightened compared to the CITED2/p300 and HIF-1α /P300 binary complexes and the C terminus of CITIF (corresponding to HIF-1α_815-826_) is not fully folded, which might be due to the allosteric effects of CITED2 residues binding.

**Figure 6.**
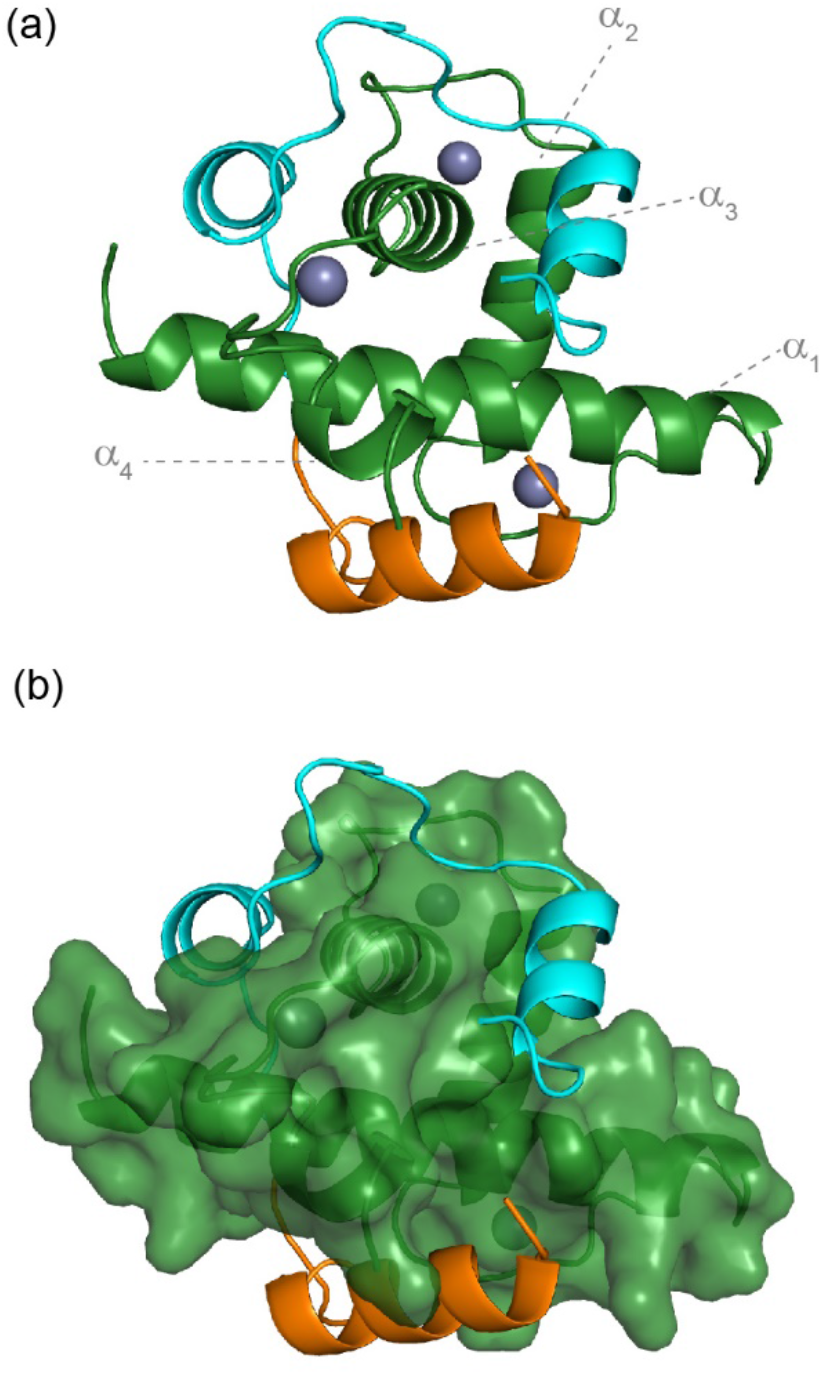
Crystal structure of p300 (green) in complex with CITIF determined at 2 Å resolution (PDB: 7QGS). Residues corresponding to CITED2_224-243_ are coloured orange, residues corresponding to HIF-1α_792-826_ are coloured cyan.

These data show that CITED2_224-243_ and HIF-1α_792-826_ sequences can bind simultaneously to p300 without interfering with one another, further supporting the hypothesis of the ternary complex formation, and suggesting that these sequences may not contain the key components that induce unidirectional displacement of HIF-1α, by CITED2. Berlow *et al*. previously used ^1^H–^15^N NOE experiments to identify significant differences in the degree of dynamic disorder and therefore flexibility between p300 bound ^15^N HIF-1α and ^15^N CITED2.^46^ HIF-1α was shown to display a wide range of dynamics throughout its sequence with both ordered and flexible regions, notably in the LPQL motif which was shown to play a role in the binding mechanism. CITED2 on the other hand elicited more uniform behaviour consistent with a more ordered structure. Subsequently Appling *et al*. used a HIF-1α-CITED2 fusion peptide similar to the one reported here to probe further the binding mechanism; these studies revealed that the region corresponding to HIF-1α_815-826_ and the region corresponding to the CITED2_224-235_ are mutually destabilizing to one another and this negative allostery is governed by the length and orientation of the C-terminal p300 helix (α_4_).^47^ Molecular dynamics simulations also identified a HIF-1α/CITED2/p300 ternary complex in support of this model and point to a role for hydrophobic residues C-terminal to the LPEL residues as being important in displacing the HIF-1α_797-805_ helix.^61-63^ Similarly, ^15^N-relaxation and side chain methyl ^2^H-relaxation experiments on p300 and side chain methyl ^2^H-relaxation for bound HIF-1α demonstrated (i) that side-chain and backbone dynamics for p300 upon binding to CTAD-HIF-1α involve an unfavourable conformational entropy change on complex formation (with the backbone contribution dominant), (ii) that HIF-1α similarly undergoes a significant side chain conformational entropy change upon p300 recognition and (iii) the N-terminal region of HIF-1α, the residues in p300 contacting the LPQL motif and the C-terminus of p300 remain dynamic when bound.^64^ Finally, comparative *in silico* alanine scanning results (Figure S10) determined using BAlaS^56^ (a web-based server version of BudeAlaScan) for HIF-1α/p300 (PDB ID: 1L8C), CITED2/p300 (PDB ID: 1P4Q) and the CITED-HIF-fusion/CBP complex (PDB ID: 7LVS) reported by Appling et al. show a dispersed distribution of potential hot residues (similar to that observed for HIF-1α/p300 (PDB ID: 1L8C) using BudeAlaScan), but with a greater proportion towards the N-terminus in CITED2 and the C-Terminus in HIF-1α. The variation of one of these hot residues (L63A) in the CITED2-HIF-1α fusion peptide corresponding to L822A in this work) resulted in the complete displacement of the C terminal HIF-1α_815-826_ helix which allowed the binding of the N terminal helix of the fusion peptide (corresponding to CITED2_216-246_).^47^ This implies that although individual variations do not have a significant effect on overall binding affinity they can be important mechanistically in mediating allosteric responses. Our ITC experiments can be rationalized in the context of these data; strongly favourable p300 binding enthalpies, and strongly unfavourable p300 binding entropies are consistent with a dynamic HIF-1α/p300 interaction, whereas the weaker enthalpies and entropies of interaction are consistent with a more ordered CITED2/p300 interaction. Crucially where CITIF is concerned, the N-terminal fragment of CITED2_224-243_ is derived from a region that is highly ordered in the CITED/p300 interaction and so the observed enthalpy-entropy compensation might be excepted for CITIF which marries the N-terminus of CITED2 with the C-Terminus of HIF-1α. Overall, the results are fully consistent with the sequence variation studies described above in which variants with a significant predicted ΔΔG were observed to bind with comparable affinity, less favourable enthalpy and more favourable entropy when compared to the parent sequence. Taken together, the results underscore recent observations on the protein-protein interactions of intrinsically disordered regions in which sequence variation has limited impact on binding affinity;^65-66^ enthalpy-entropy compensation provides the scope for such fuzzy interactions to accommodate sequence variation without significant impact on binding affinity and therefore function.

## Conclusions

We have shown using a combination of single and multiple alanine sequence variants of HIF-1α alongside sequence hybrids with the negative regulator of HIF-1α (CITED2) that interaction with p300 is highly tolerant to sequence variation as demonstrated by fluorescence anisotropy and isothermal titration calorimetry. Recent studies on the interaction of p300(CBP) with HIF-1α or CITED2 have largely focussed on dynamic structural studies and molecular dynamics simulations to rationalise the displacement of HIF-1α from p300 by CITED2.^40, 46-47, 53, 61-64^ Our equilibrium measurements for a range of sequence variants provide complementary data demonstrating interaction between HIF-1α and p300 is characterized by a large favourable enthalpy and large unfavourable entropy of binding. The absence of dramatic changes in binding affinity for alanine variants taken together with an observed enthalpy-entropy compensation is consistent with significant chelate co-operativity^21, 50^ and dispersal of binding energy across the sequence, with binding free energy derived from non-covalent contacts between the HIF-1α backbone (in addition to side-chains) and the surface of the p300 CH1 domain, alongside favourable long range electrostatic and transient side-chain interactions during HIF-1α folding. Such behaviour provides a mechanism for the intrinsically disordered HIF-1α sequence to tolerate sequence variation by adopting a more disordered bound state in its interaction with p300. Binding of CITED2 to p300 however is characterized by small favourable enthalpy and entropy changes, yet (in our hands) its affinity for p300 is slightly higher than that of HIF-1α and therefore may also augment the allosteric changes that accompany ternary complex formation between HIF-1α, CITED2 and p300 *en route* to unidirectional displacement of HIF-1α by CITED2. Such behaviour is encompassed in CITIF, a HIF-1α-CITED2 hybrid sequence; p300 affinity is higher than HIF-1α and comparable to CITED2, with a thermodynamic signature that is intermediate between the two representing a consonance between the high affinity less dynamic CITED2 sequence and the more fuzzy HIF-1α. This and the sequence dependent competition mechanism by which the negative feedback regulator CITED2 displaces HIF-1α may provide insight to inform design of selective HIF-1α modulators. More broadly, these results underscore the advantageous features of intrinsically disordered regions in facilitating function^67-68^ whilst such sequence tolerance may represent an additional rational for the prevalence of disease relevant mutations within intrinsically disordered regions.^69^

## Abbreviations list

ALA: scan Alanine scan
BUDE: Bristol University Docking Engine
C-TAD: Carboxy-terminal transactivation domain
FA: Fluorescence Anisotropy
GFP: Green Fluorescent Protein
GST: Glutathione S-transferase
HIF-1α: Hypoxia-inducible factor 1-alpha
ITC: Isothermal titration calorimetry
mAV: multiple alanine variant
NMR: Nuclear magnetic resonance
PPI: Protein-protein interaction
sAV: Single alanine variant
wt: wild type

## Declarations of interest

The authors declare no competing financial interests.

## Acknowledgements

We would like to thank Dr Iain Manfield for his support with ITC measurements and ongoing collaboration with AstraZeneca, Domainex and the Northern Institute for Cancer Research. We would like to thank Dr Chi H. Trinh for his support with crystallography. We acknowledge Diamond Light Source for time on Beamline I04 under Proposal mx19248 and the authors would like to thank the staff of this beamline for assistance with data collection.

## Funding information

This work was supported by EPSRC (EP/N013573/1 and EP/KO39292/1) and the BBSRC/EPSRC-funded Synthetic Biology Research Centre, BrisSynBio (BB/L01386X/1). This project has received funding from the European Union’s Horizon 2020 research and innovation programme under the Marie Skłodowska-Curie grant agreement no. MSCA-IF-2016-749012. Z.H. received funding from the National Research, Development and Innovation Office – NKFIH PD 135324. A. J. W. wishes to acknowledge the support of a Royal Society Leverhulme Trust Senior Fellowship (SRF\R1\191087)

## Author contribution statement

Z. H., A.J.W. and T. A. E. conceived and designed the research program, F. H and Z. H. designed studies and performed the research with support from A. A. I., V. L. P., G. J. B, and R. B. S. The manuscript was written by F. H, and Z. H. and edited into its final form by A.J.W. and T.A.E with contributions from all authors.

## Notes

### Competing Interest Statement

The authors have declared no competing interest.

