## Supplemental Information for "Understanding p300-transcription factor interactions using sequence variation and hybridization"

### Supporting information

##### Abstract

The hypoxic response is central to cell function and plays a significant role in the growth and survival of solid tumours. HIF-1 regulates the hypoxic response by activating over 100 genes responsible for adaptation to hypoxia, making it a potential target for anticancer drug discovery. Although there is significant structural and mechanistic understanding of the interaction between HIF-1α and p300 alongside negative regulators of HIF-1α such as CITED2, there remains a need to further understand the sequence determinants of binding. In this work we use a combination of protein expression, chemical synthesis, fluorescence anisotropy and isothermal titration calorimetry for HIF-1α sequence variants and a HIF-1α-CITED hybrid sequence which we term CITIF. We show the HIF-1α sequence is highly tolerant to sequence variation through reduced enthalpic and less unfavourable entropic contributions, These data imply backbone as opposed to side chain interactions and ligand folding control the binding interaction and that sequence variations are tolerated as a result of adopting a more disordered bound interaction or "fuzzy" complex.

#### Supplementary Figures and Tables

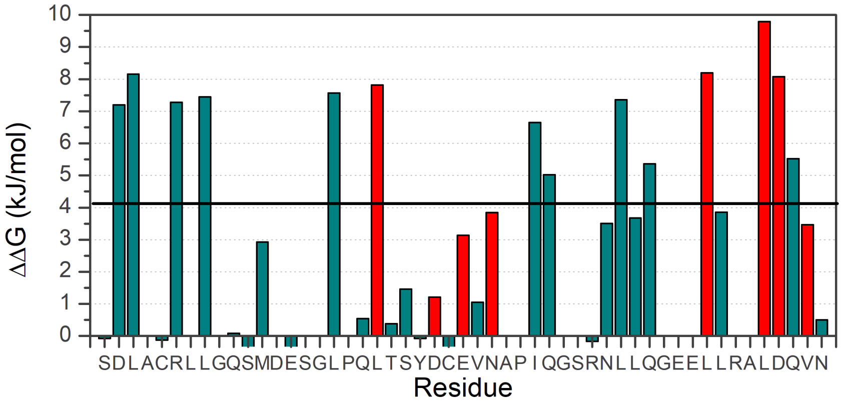

**Figure S1** Binding energy changes predicted by Robetta for the HIF-1α/p300 interaction using the lowest energy structure from the NMR derived ensemble PDB ID: 1L8C.

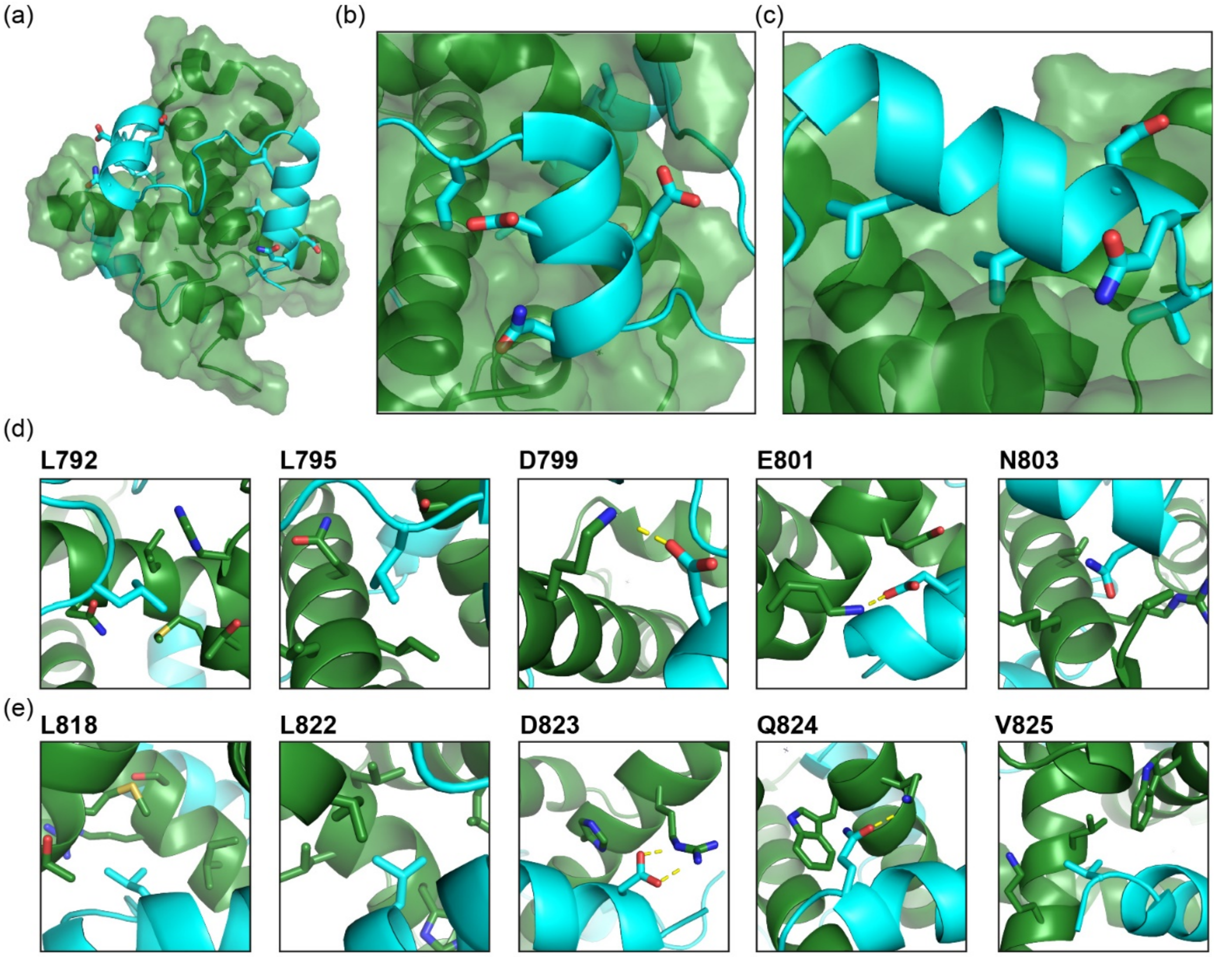

**Figure S2** Structural visualization of predicted hot residues. (a) surface representation of p300 (green) bound to HIF-1α (cyan ribbon) highlighting the C-terminal helices (HIF-1α$_{797-805}$ and HIF-1α$_{815-826}$); (b) hot residues found in the HIF-1α$_{797-805}$ helix; (c) hot residues found in the HIF-1α$_{815-826}$ helix; (d) intermolecular interactions of individual hot-residues from HIF-1α$_{797-805}$ helix (e) intermolecular interactions of individual hot-residues from HIF-1α$_{815-826}$ helix (HIF-1α is shown in cyan, p300 in green and selected hotspots are shown as stick representation, polar contacts indicated with yellow dashed lines)

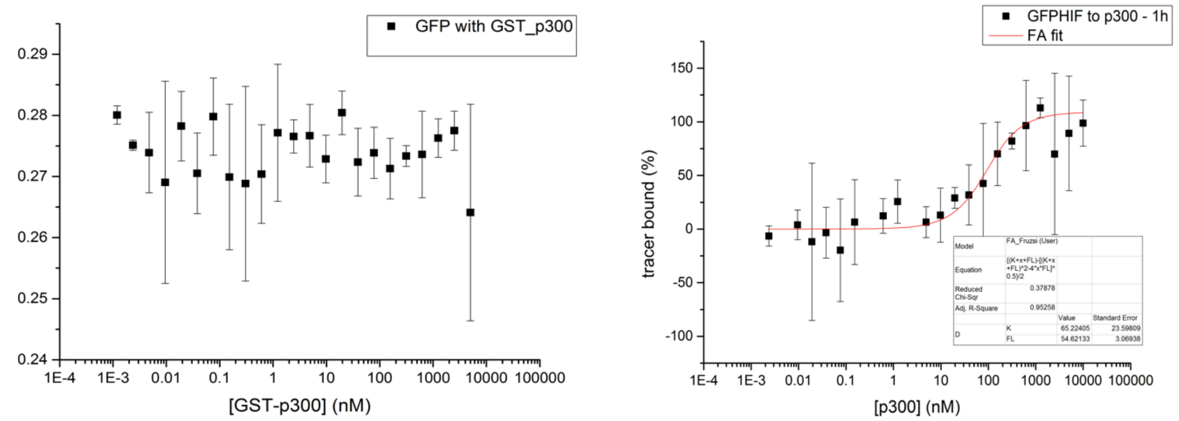

**Figure S3**Fluorescence anisotropy direct titrations of 50nM GFP with increasing concentration of GST-p300 (top) and GFP-HIF-1α$_{776\text{-}826}$ with p300 to show that GFP does not interact with GST-p300 nor does the presence of GST affect the binding to GFP-HIF-1α. (25 mM Tris-HCl, 150 mM NaCl, 1mM DTT, pH 7.4). The data was analysed using Origin, highest concentration of p300 was 10μM, with a two times dilution series applied.

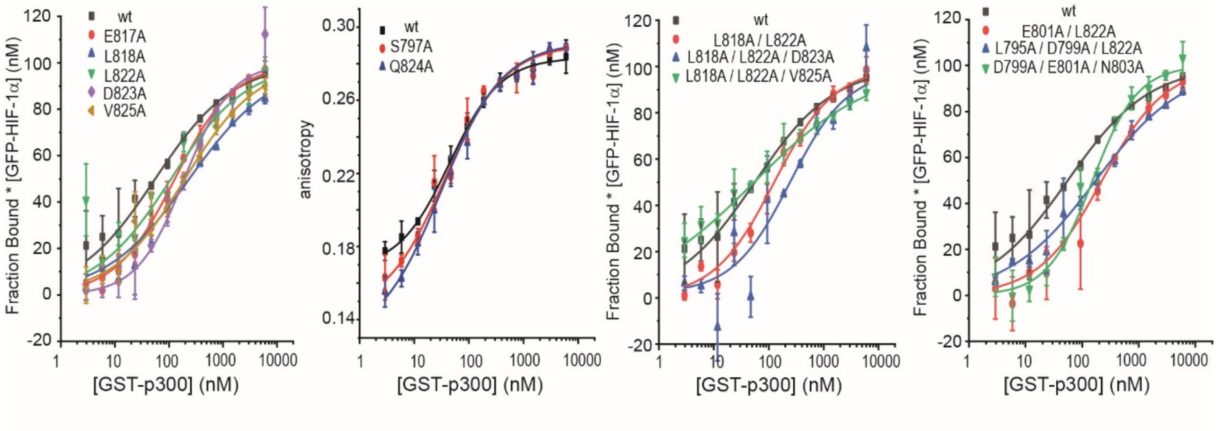

**Figure S4**Representative fluorescence anisotropy titration data for sAV and mAV HIF-1α peptides interacting with p300 (25 mM Tris-HCl, 150 mM NaCl, 1mM DTT, pH 7.4);

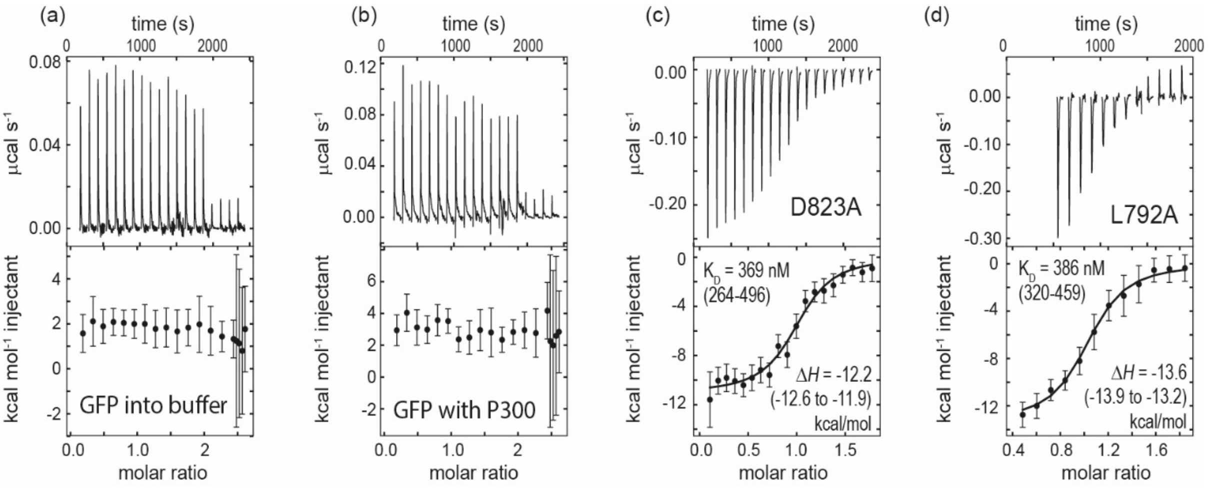

**Figure S5** Heat of dilution of (a) GFP titrated to buffer, (b) GFP titrated to p300 and (c) Raw ITC (upper) data and fitted thermogram (lower) for the interaction of GFP-HIF-1α$_{776\text{-}826}$ D823A variant (37$^{o}$C in 25 mM Tris-HCl, 150 mM NaCl, 1mM DTT, pH 7.4) using 10 μM p300 in the cell and 100 μM HIF-1α variant in the syringe. ITC experiments were carried out using a Microcal ITC200i instrument (Malvern). All proteins were dialysed against the same buffer prior to experiment, buffer or 10 μM p300 was present in the cell and titrated with 100 μM GFP-HIF-1α or GFP loaded into the syringe using 16 x 2 μL injections with 120 s spacing between the injections. Heats of GFP or GFP-HIF-1α dilution were subtracted from the measurement raw data.

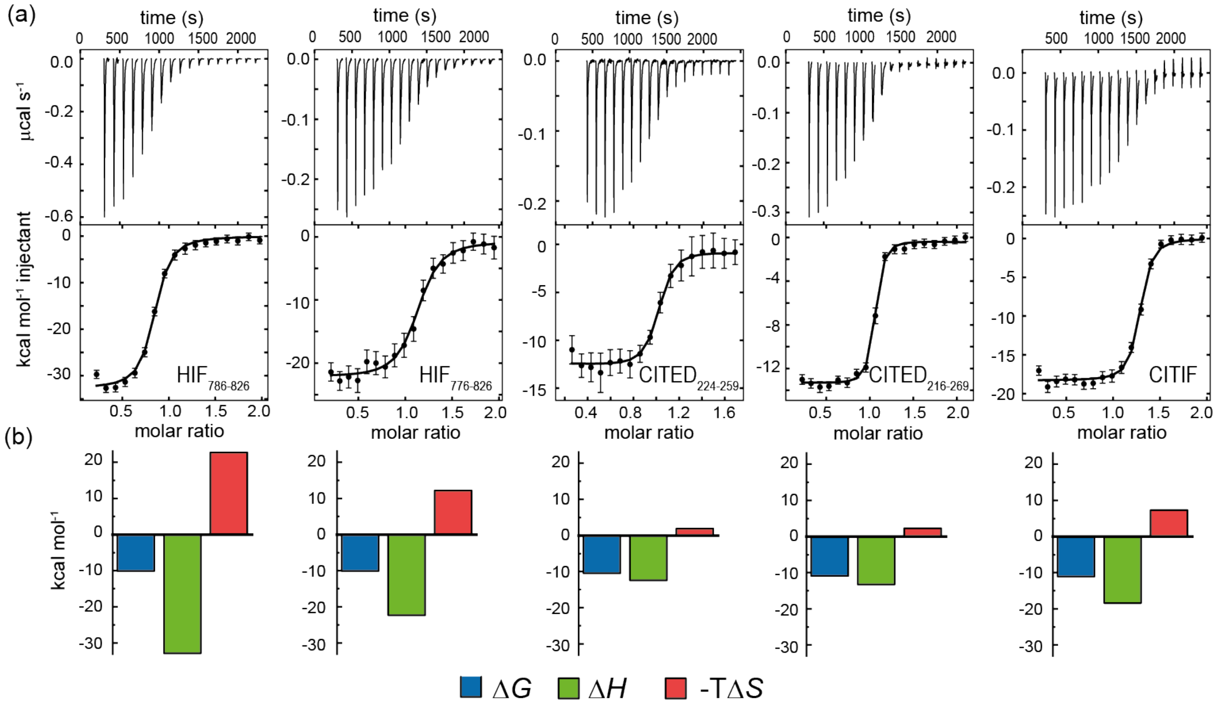

**Figure S6** Isothermal titration calorimetry data for the interaction of expressed HIF-1α$_{776\text{-}826}$, HIF-1α$_{786\text{-}826}$ CITED2$_{224\text{-}259}$, CITED2$_{216\text{-}269}$ and CITIF peptides with p300. (a) Raw ITC (upper) data and fitted thermogram (lower) (40 mM sodium phosphate, pH 7.5 100 mM NaCl, 1 mM DTT buffer using 5 µM protein in the cell and 60 µM ligand in the syringe at 35°C); (b) Thermodynamic signatures for each interaction.

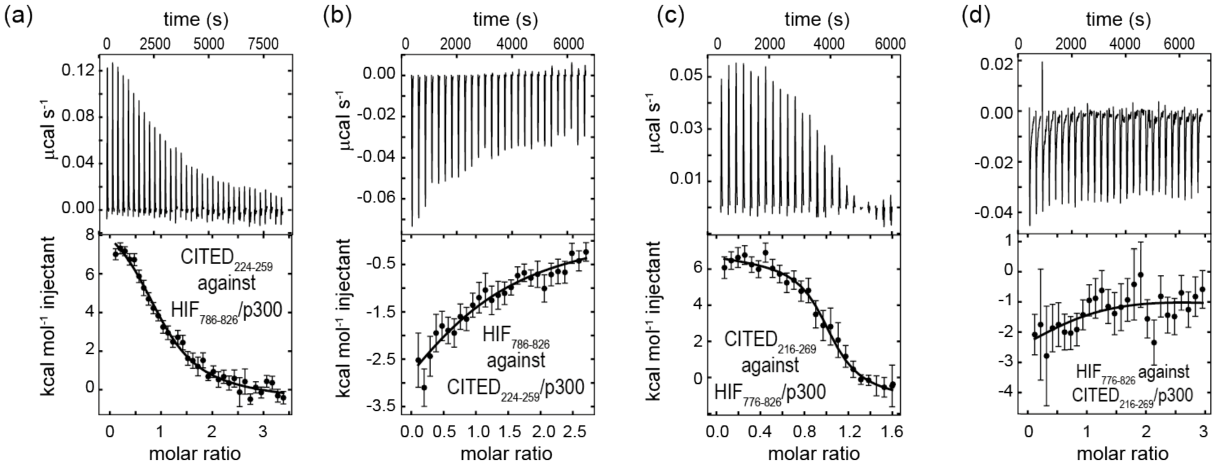

**Figure S7** Apparent $K_D$ fits for the competition ITC experiment using a binary interaction model. p300/ligand complexes were prepared by titrating the appropriate ligand first to p300 in the cell using 3-5 µM protein concentration, the titration stopped when it reached close to saturation conditions, which resulted in 1.15-1.5 excess of ligand compared to p300 in the cell (Table S4) Then it was titrated with the competitor ligand using 30-60 µM concentration in the syringe, 5 or 10 uL injections with 180-240 s spacing. Background titrations were performed without p300 in the cell and subtracted from the measurements. Apparent $K_{D,app}$ and $\Delta H_{app}$ values data were fitted to a binary interaction model in SEDPHAT.

**Table S1** Estimated and fitted apparent $K_D$ values and thermodynamic parameters for competition ITC experiments. Estimated parameters are based on maximum negative cooperativity (α =0, Δ$\Delta H$ =0, see equations in experimental section and Table S4 for concentrations) Data were fitted using a binary interaction model to give apparent $K_D$ and $\Delta H$ values, 68% confidence intervals are shown in brackets.

| | ***Estimated parameters*** | | ***Measured binding parameters*** | | |
|---|---|---|---|---|---|
| | $K_{D,app}$ µM | $\Delta H_{app}$ kcal/mol | $K_{D,app}$ µM | $\Delta H_{app}$ kcal/mol | $\Delta S_{app}$ cal/mol/K |
| **CITED$_{224-259}$ titrated to HIF$_{786-826}$/P300 complex** | 1.2 | 13.2 | 0.665 (0.840, 0.490) | 9.46 (8.96, 10.00) | 58.9 |
| **HIF$_{786-826}$ titrated to CITED$_{224-259}$ /P300 complex** | 26 | -13.2 | 5.0 (6.50, 2.9) | -6.6 (-7.4, -6.0) | 2.6 |
| **CITED$_{216-269}$ titrated to HIF$_{776-826}$/P300 complex** | **1.7** | 8.7 | **0.043** (0.053, 0.034) | 6.76 (6.59, 6.90) | 55.6 |
| **HIF$_{776-826}$ titrated to CITED$_{216-259}$ /P300 complex** | 12 | -9.2 | 2.4 (6, 0.9) | -4.2 (-6.2, -3.1) | 5.8 |
| **CITED$_{216-248}$ titrated to HIF$_{776-826}$/P300 complex** | **35** | 17 | **2.0** (1.8, 2.2) | 7.9 (7.4 – 8.4) | 51.7 |

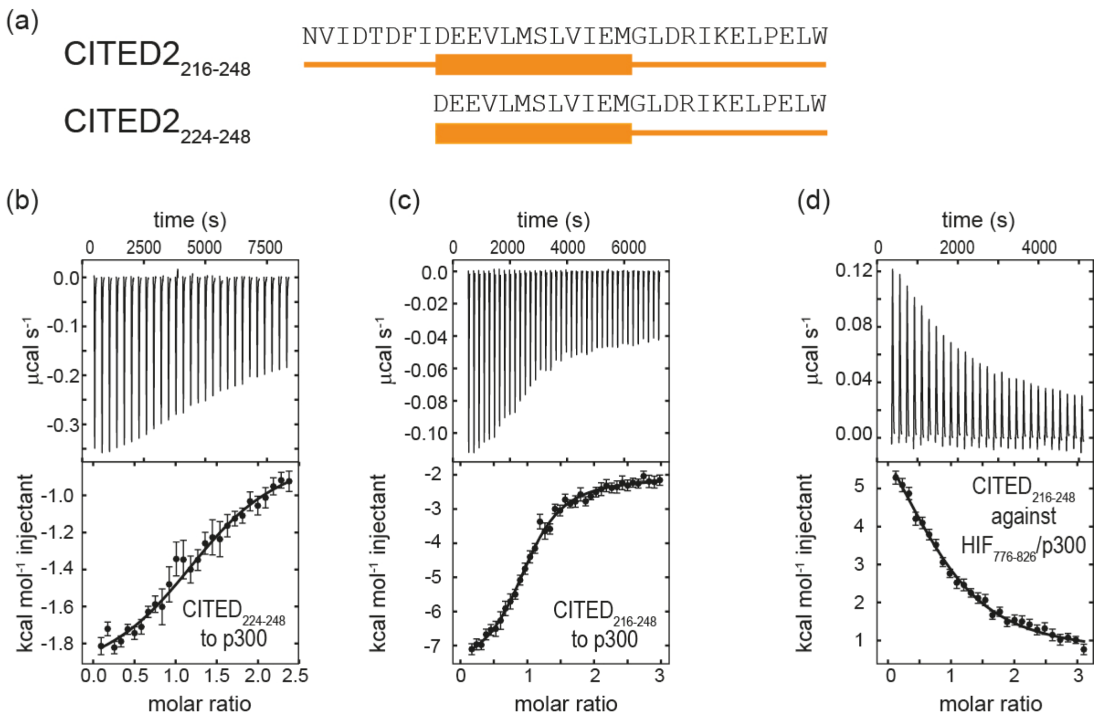

**Figure S8** (a) Sequences of the truncated CITED2 peptides and isothermal titration calorimetry data for (b) $CITED_{224\text{-}248}$ titrated to p300 using 45 μM p300 in the cell and 500 μM peptide in the syringe and (c) $CITED_{216\text{-}248}$ titrated to p300 using 5 μM p300 in the cell and 50 μM peptide in the syringe. (d) Competition ITC measurement for displacing $HIF_{776\text{-}826}$ by $CITED_{216\text{-}248}$ (conditions listed in Table S4). All measurements were carried out 40 mM sodium phosphate, pH 7.5 100 mM NaCl, 1 mM DTT buffer at 35°C.

**Table S2** Thermodynamic parameters for shortened CITED2 sequences. 68% confidence intervals are shown in brackets.

| | | $K_D$, μM | $\Delta H$ kcal mol$^{-1}$ | $\Delta S$ cal mol$^{-1}$ K$^{-1}$ | $\Delta\Delta G$ kcal mol$^{-1}$ | $\Delta\Delta H$ kcal mol$^{-1}$ |
|---|---|---|---|---|---|---|
| direct titration to p300[a] | **$CITED2_{224\text{-}248}$** | 10.44 (7.86 – 14.2) | -1.28 (-1.45 to -1.15) | 18.62 | | |
| | **$CITED2_{216\text{-}248}$** | 0.303 (0.230 – 0.397) | -5.7 (-6.1 to -5.4) | 11.3 | | |
| $CITED2_{216\text{-}248}$ titrated to $HIF_{776\text{-}826}$/p300 complex[b] | **$CITED2_{216\text{-}248}$** | 0.297 (0.238 – 0.373) | -5.7 (-5.9 to -5.4) | 11.3 | 1.27 (0.9 – 1.69) | 9.8 (8.5 – 11.3) |
| | **$HIF_{776\text{-}826}$** | 0.067 (0.059 – 0.075) | -27.8 (-28.1 to -27.5) | -57.4 | | |

[a] Data fitted to a single binding site model

[b] Data fitted using global fitting method, using a cooperative binding model including a ternary complex formation and a fit for ΔΔG and ΔΔH.

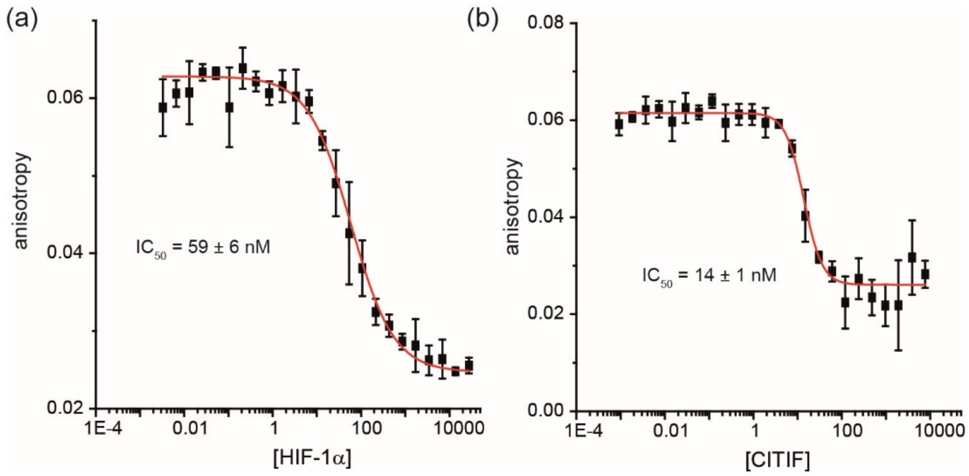

**Figure S9** Fluorescence anisotropy competition titration data for: (a) HIF-1α$_{786-826}$ and (b) CITIF peptides illustrating that CITIF competes with HIF-1α for p300 (25 mM Tris-HCl, 150 mM NaCl, 1mM DTT, pH 7.4 using 5 nM FAM-HIF$_{786-826}$ as tracer, with a half dilution series of p300 from starting from 10 μM over 24 points); (note: whilst the FA competition experiments for the expressed peptides revealed some differences in IC$_{50}$ values that are consistent with a higher affinity of CITIF than HIF-1α for p300 the concentrations of HIF-1α and p300 in the assay are such that inhibition occurs at the limit of the assay, thus these numbers should be treated with caution).

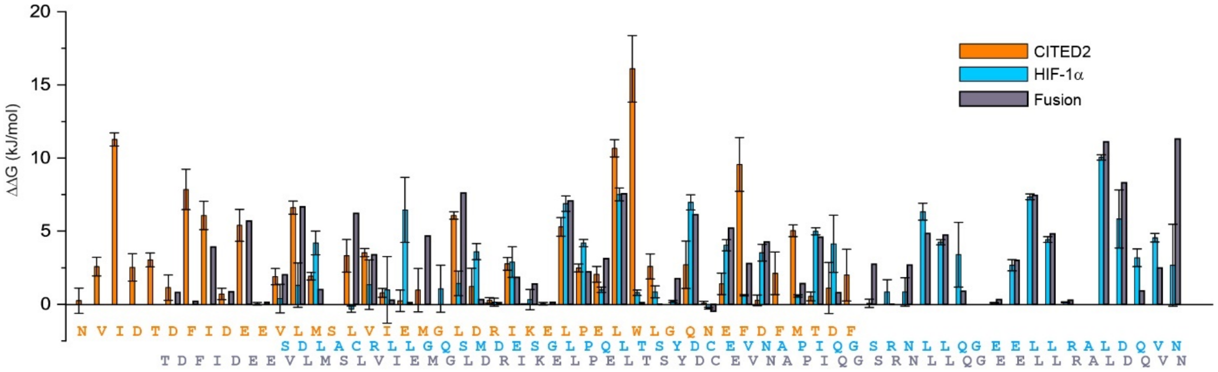

**Figure S10** *in silico* Alanine scanning data were calculated for HIF-1α/p300 (PDB ID: 1L8C), CITED2/p300 (PDB ID: 1P4Q) and CITED-HIF-fusion/CBP (PDB ID: 7LVS) structures using the BAlaS server (https://pragmaticproteindesign.bio.ed.ac.uk/balas/).[1]

**Table S3** Statistics obtained for the p300/CITIF co-crystal structure

| | |
|---|---|
| **Reservoir conditions** | 0.1 M HEPES pH 6.5, PEG 6K 35% |
| **Data Collection** | |
| X-ray source | DLS Beamline i04 |
| Processed using | xia2 (DIALS, Aimless) |
| oscillations | 0.1 |
| images collected | 3600 |
| Space group | $P\ 1\ 2_1\ 1$ |
| **Unit cell dimensions** | |
| *a*, *b*, *c*, (Å) | 30.50, 48.60, 39.83 |
| $\alpha$, $\beta$, $\gamma$ (°) | 90, 103.87, 90 |
| Resolution | 2.0 - 29.6 (2.0 - 2.07) |
| Observations | 51028 (2141) |
| Unique reflections | 7688 (747) |
| $R_{merge}$ (I) | 0.109 (0.744) |
| $R_{meas}$ (I) | 0.119 (0.814) |
| $R_{pim}$ (I) | 0.047 (0.325) |
| CC 1/2 | 0.991 (0.870) |
| I/$\sigma$ | 7.5 (2.9) |
| Completeness | 99.7 (96.5) |
| Redundancy | 6.6 (6.0) |
| **Refinement** | |
| Protein molecules in au | 2 |
| Rwork/Rfree | 0.221/0.245 |
| No atoms | 1210 |
| Protein | 1166 |
| Ligand | 3 (Zn) |
| Water | 41 |
| **Mean B factors (Å)** | |
| Protein | 79.14 |
| Ligand | 57.35 |
| Water | 77.69 |
| **R.m.s. deviations** | |
| Bond length (Å) | 0.016 |
| Bond angles | 1.55 |
| **Ramachandran statistics** | |
| % favoured | 95.14 |
| % allowed | 4.86 |
| % outliers | 0 |
| Clashscore | 13.3 |

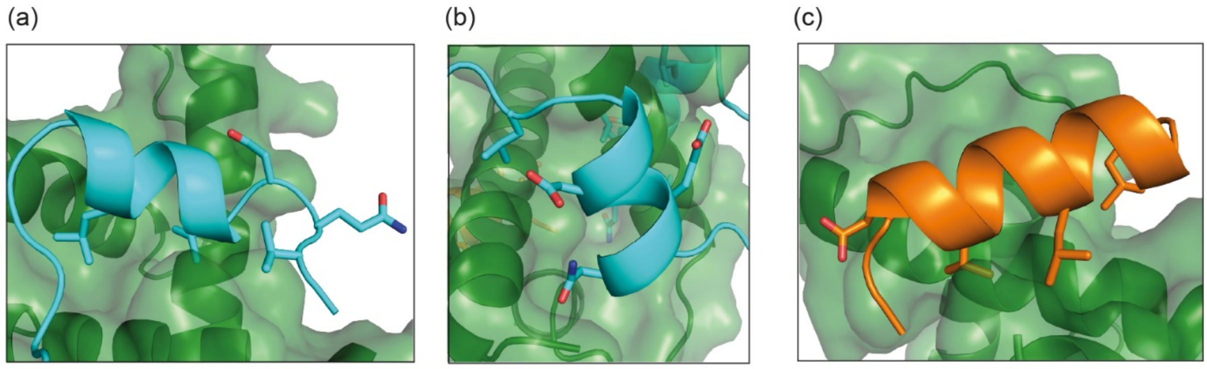

**Figure S11** Helical regions of the CITIF/p300 complex crystal structure (PDB: 7QGS), hot residues are shown as stick representation. (a) Residues corresponding to HIF-1α$_{815-826}$ helix (b) residues corresponding to HIF-1α$_{797-805}$ helix (c) residues corresponding to helix CITED2$_{224-237}$ helix.

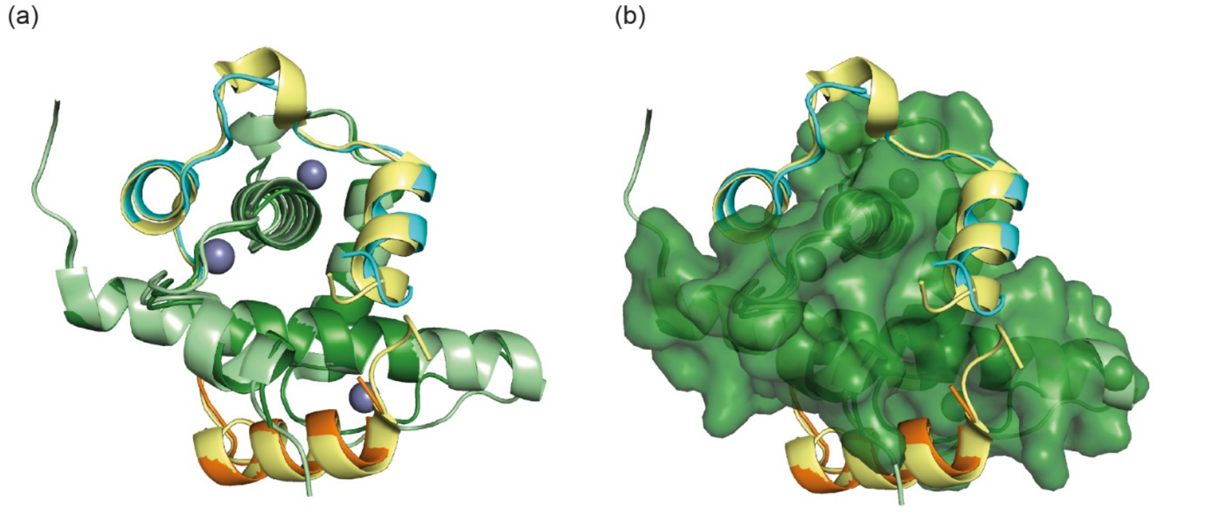

**Figure S12** Overlay of the CITED2-HIF-1α fusion peptide/CBP crystal structure (PDB:7LVS, CBP in light green, fusion peptide in yellow) with the CITIF/p300 crystal structure (PDB:7QGS) p300 in green, CITIF in cyan/orange (cyan corresponding to HIF-1α$_{792-826}$ and orange corresponding to CITED2$_{224-243}$ residues). Only minor differences are observed between the two structures, with the only notable conformational difference observed for Gln39-Leu44 (7QGS numbering), whereby a more linear conformation is adopted in 7QGS.

### Materials and Methods

#### *In silico* predictions

##### Alanine scan using Robetta

Alanine scanning data for HIF-1α/p300 were calculated with the Robetta server, http://robetta.bakerlab.org, using the lowest energy structure from the NMR derived ensemble PDB ID: 1L8C.

##### BUDE Alanine scan

Alanine scanning data (show in Fig. 1g) for HIF-1α /p300 were calculated using BudeAlaScan as previously described[1] using the NMR derived ensemble PDB ID: 1L8C (standard deviations represent the variation in determined ΔΔG between individual structures in from the ensemble). Additionally alanine scanning data were calculated for the HIF-1α/p300, CITED2/p300 and CITED-HIF-fusion/CBP structures using PDB ID: 1L8C, 1p4q and 7lvs respectively using the BalaS server (https://pragmaticproteindesign.bio.ed.ac.uk/balas/).[1]

#### Plasmids for protein production

The DNA sequence of human p300 (UniProtKB - Q09472) containing the CH1 domain (Zinc finger, TAZ type 1) from residues 330-420 was cloned into pGEX-6P-2 or pGEX-4T-1 plasmids using BamHI/XhoI restriction sites.

The genes encoding *H sapiens* HIF-1α (residues 776-826 and 786-826, Uniprot Q16665), CITED2 (residues 224-255 and 216-269, Uniprot Q99967) and CITIF (containing CITED2 224-243 fused with HIF-1α residues 792-826) were inserted into pET28:GFP plasmid using BamHI/XhoI restriction sites creating $His_{10}$-GFP constructs. HIF-1α mutants (residues 776-826) were produced using the Quikchange Site Directed Mutagenesis Kit (Agilent) and the wild type protein expression vector as template (oligo sequences are shown in Table S5). The sequences of all constructs were confirmed by DNA sequencing prior to expression.

#### Expression and purification of p300 CH1 domain

Expression and purification of GST-p300 was performed as described previously[2] using either pGEX-6P2 or pGEX-4T1 plasmids. *E. coli* strain Rosetta 2 (transformed with pGEX-6P2-p300) or BL21 E.Coli pLysS (transformed with pGEX-4T1-p300) were grown at 37°C to OD600 0.6-0.8 and induced with 0.1 mM IPTG and 50 µM $ZnSO_4$ was added and incubated overnight at

18 °C. Cells were harvested and lysed by sonication and centrifuged at 25.000 g for 30 minutes at 4ºC. The supernatant was applied to glutathione beads (Glutathione Sepharose 4B, GE Healthcare) and washed with 20 mM Tris, pH 8, 150 mM NaCl and 20 mM Tris, pH 8, 500 mM NaCl. GST was cleaved on-column overnight at 4ºC using PreScission protease (pGEX-6P-2 plasmid) or Thrombin protease (pGEX-4T-1 plasmid). For the fluorescence anisotropy (FA) experiments the GST tag cleavage step was omitted and the GST tagged protein was eluted using 20 mM glutathione containing buffer. The eluted fractions were concentrated and purified by size-exclusion chromatography on S75 26/60 pg column in 20 mM Tris, 150 mM NaCl, 5% glycerol, 1mM DTT pH 7.5 buffer. Collected fractions were analysed by SDS-PAGE and pure fractions were concentrated. Pure protein was analysed by high resolution mass spectrometry: For the PreScission cleaved construct: expected m/z: 10870.63 measured m/z: 10864.2471. This construct was used to perform ITC experiments on MicroCal iTC200 instrument (titrated with expressed interaction partners). For thrombin cleaved construct expected m/z: 10603.30, measured m/z: 10603.41, this construct was used to perform experiments on Microcal VP ITC (titrated with synthetic interaction partners). Concentration was determined by UV-VIS spectroscopy using 5500 $M^{-1}$ $cm^{-1}$ extinction coefficient.

**a** GPIGSA$_{330}$DPEKRKLIQQQLVLLLHAHKCQRREQ
ANGEVRQCNLPHCRTMKNVLNHMTHCQSGKSCQV
AHCASSRQIISHWKNCTRHDCPVCLPLKNA$_{420}$

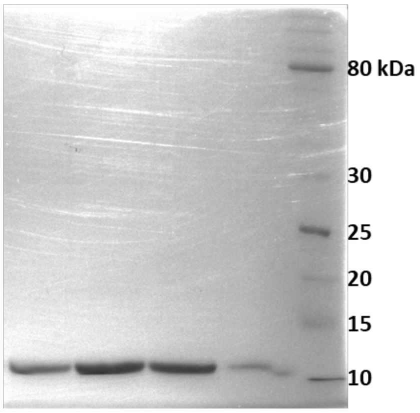

**b** GSA$_{330}$DPEKRKLIQQQLVLLLHAHKCQRREQAN
GEVRQCNLPHCRTMKNVLNHMTHCQSGKSCQVA
HCASSRQIISHWKNCTRHDCPVCLPLKNA$_{420}$

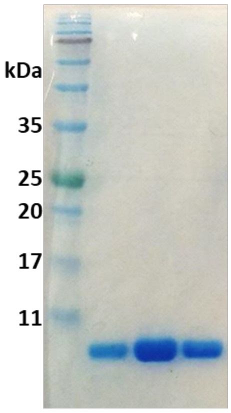

**Figure S13** Sequence and SDS gel of purified p300 CH1 using a) pGEX-6P-2 plasmid followed by PreScission protease cleavage and b) using pGEX-4T-1 plasmid and thrombin cleavage.

#### Expression and purification of GFP tagged HIF-1α and GFP-HIF-1α alanine variants

GFP-HIF-1α residues 776-826 and its alanine variants were expressed in the *E. coli* strain Rosetta 2. Bacterial cultures were grown at 37°C to a density of $OD_{600}$ = 0.5-0.8 and protein expression then induced by addition of 1 mM isopropyl-β-D- thiogalactoside (IPTG). Upon induction, cultures were cooled and maintained at 22°C for protein expression overnight. Cell pellets were harvested by centrifugation for 7 minutes at 8655 x g and then re-suspended in lysis buffer (25 mM Tris-HCl pH 8, 500 mM NaCl) containing (1 mg $mL^{-1}$ lysozyme, 10mg $mL^{-1}$ of DNase1 and 1 tablet per 1 L culture protease inhibitor tablets (Complete Mini, Roche) and lysed by sonication. The cell lysate was then centrifuged for 45 minutes at 23655 x g, the supernatant was collected and applied onto a Ni-Sepharose resin column (bed size 5 mL). The column was washed with 40 mL wash buffer I (25 mM Tris-HCl, 500 mM NaCl, pH 8), 40 mL wash buffer II (25 mM Tris-HCl, 15 mM Imidazole, 500 mM NaCl, pH 8) and 400mL was buffer III (25 mM Tris-HCl, 50 mM Imidazole, 500 mM NaCl, pH 8). Bound proteins were eluted with 10 mL of elution buffer (25 mM Tris-HCl, 400 mM Imidazole, 500 mM NaCl, pH 8). The elution fraction was loaded on Superdex75 (26/600) prep grade size exclusion chromatography column, equilibrated with Gel filtration buffer (25 mM Tris-HCl, 150 mM NaCl, 1mM DTT, pH 7.4). Samples of the fractionated eluent were applied to 15% SDS gels to assess sample purity and the purified protein was concentrated using Amicon-15 10K (Merck) centrifugal concentrators with a MWCO of 10kD and stored at -80°C in small aliquots.

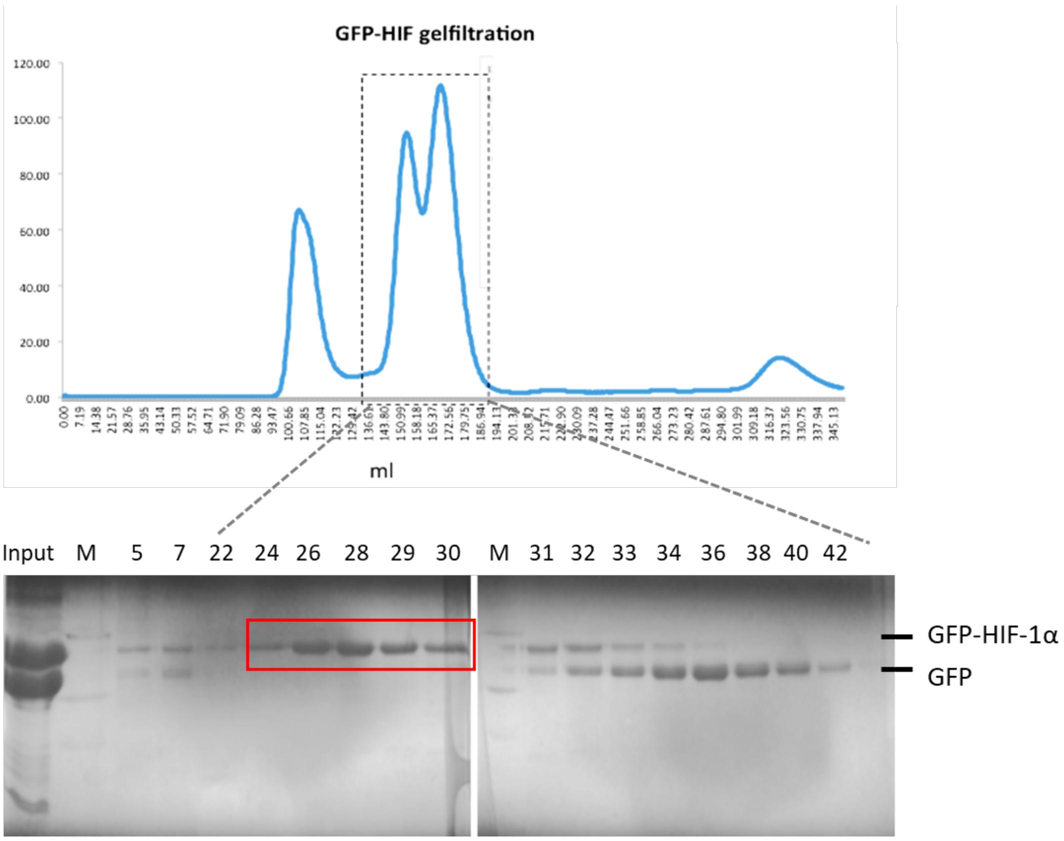

**Figure S14** Purification of recombinantly expressed GFP-HIF-1α$_{776\text{-}826}$ by size exclusion chromatography. During the initial purification step the fusion-protein is partially cleaved, but the cleaved GFP could be separated from GFP-HIF-1α by size-exclusion chromatography (red square), and later the protein remained stable.

#### Expression and purification of untagged HIF-1α, CITED2 and CITIF

Untagged HIF-1α (residues 776-826 and 786-826), CITED2 (residues 224-255 and 216-269) and CITIF were expressed using the same conditions as described above with the following modifications. After cell lysis the inclusion bodies were washed with lysis buffer and the insoluble pellet was resuspended in 6M urea, 0.5 M NaCl, 5 mM imidazole, 50 mM Tris pH 8, 1 mM DTT and sonicated and centrifuged as before. The supernatant was applied to a His-trap HP column (5ml) and washed with 10 column volumes of the resuspension buffer, then with 4 M urea, 0.5 M NaCl, 20 mM imidazole, 20 mM Tris, pH 8, 1 mM DTT, followed by washing with 2 M urea, 0.25 M NaCl, 20 mM imidazole, 20 mM Tris, pH 8, 1 mM DTT then 20 mM Tris, 0.1 M NaCl, pH 8, 1mM DTT. PreScission protease was added to the last buffer and 5 ml applied to the column and on-column cleavage performed at 4°C. Cleavage was followed by HRMS. The eluted protein without further concentration, was immediately purified using

RP-HPLC on an Agilent 1260 Infinity HPLC equipped with a diode array detector using a Kinetex EVO C18 column. Eluent A: H2O + 0.1% TFA, B: ACN + 0.1 % TFA. Gradient timetable: 0 min 5% B, 10 min 45% B, 25 min: 75% B, at a 10 ml/min flow rate. The collected fractions were freeze dried and analysed using high resolution mass-spectrometry (See peptide characterization data.)

##### Peptide synthesis and purification

Fmoc amino acids, Diisopropylcarbodiimide and Oxyma (Ethyl cyano(hydroxyimino)acetate) were purchased from Fluorochem. DMF (HPLC grade) and ACN (HPLC grade) was purchased form VWR, piperazine from Molar Chemicals.

Peptides were synthesized using a microwave assisted automated peptide synthesizer (CEM Liberty Blue). Preparative RP-HPLC was carried out on JASCO PU-4180 system equipped with a diode array detector (MD-4015) and an automatic fraction collector (Advantec, CHF122SC). Fraction collection was monitored and programmed using ChromNAV software. HPLC-MS measurements were carried out on a Dionex Ultimate 3000 HPLC system equipped with a diode array detector and interfaced to an LTQ XL (Thermo Scientific) ion trap mass spectrometer. Analytical UV measurements were carried out on a Shimadzu LC-20AD equipped with a SPD20A UV detector. Eluents for HPLC-UV: A: 0.1% TFA/$H_2O$; B: 0.1% TFA/ACN. Eluents for HPLC-MS: A: 0.1% HCOOH/$H_2O$; B: 0.1% HCOOH/ACN.

Peptides were synthesized on a 0.1 mmol scale on Tentagel R RAM resin (resin loading 0.19 mmol/g, Iris Biotech). Couplings were performed using 5 equivalent amino acid excesses using DIC and Oxyma as coupling reagents, dissolved in DMF. Required amounts were calculated using the built-in reagent calculator of Liberty Blue. All amino acids were double coupled using high swelling (HS) Liberty Blue methods. Deprotection solution contained 10% (w/v) piperazine dissolved in 10% absolute ethanol/NMP mixture, which effectively inhibited aspartimide formation.

###### Cycles for automated peptide synthesis

*Resin swelling* (HS): DMF (15 mL), 5 minutes

*Coupling cycle*

Deprotection: 4mL, Microwave method: Standard Deprotection (Bubble: 2s on, 3s off; 75°C 175 W 15s, 90°C 30W 50s)

4x Wash: 4 mL

2x Coupling: Amino acid 2.5 mL, Activator 1 mL, Activator Base 0.5 mL, manifold wash volume 2 mL. Microwave method: Standard coupling (Bubble 2s on, 3s off; 75 °C 183 W 15s; 90°C 30W 110s)

2x Wash: 4 mL

*Arginine coupling*

Deprotection: 4mL, 75°C Initial Deprotection (Bubble 2s on, 3s off; 75°C 40W 30s)

Deprotection: Microwave method: Standard Deprotection (Bubble: 2s on, 3s off; 75°C 175 W 15s, 90°C 30W 50s)

4x Wash: 4 mL

Coupling: Amino acid 2.5 mL, Activator 1 mL, Activator Base 0.5 mL, manifold wash volume 2 mL. Microwave method: Bubble 2s on, 3s off; 25 °C 0 W 1500s; 75°C 30W 120s

Wash: 4 mL

Coupling: 75°C Coupling, Bubble 2s on, 3s off; 75°C 30W 300s

2x Wash: 4 mL

*Cysteine coupling*

Deprotection: 4mL, 75°C Initial Deprotection (Bubble 2s on, 3s off; 75°C 40W 30s)

Deprotection: 75°C Deprotection, Bubble 2s on, 3s off; 75°C 40W 180s

4x Wash: 4 mL

Coupling: Amino acid 2.5 mL, Activator 1 mL, Activator Base 0.5 mL, manifold wash volume 2 mL. Microwave method: 50°C coupling: 25°C 0W 120 s; 50°C 35W 240s

Coupling: Amino acid 2.5 mL, Activator 1 mL, Activator Base 0.5 mL, manifold wash volume 2 mL. Microwave method: 50°C coupling: 25°C 0W 120 s; 50°C 35W 240s

2x Wash: 4 mL

*Final deprotection*

Deprotection: 4 mL, Microwave method: Standard Deprotection (Bubble: 2s on, 3s off; 75°C 175 W 15s, 90°C 30W 50s)

4x Wash

**Acetylation**

Peptides were acetylated using 10 equivalents of acetic anhydride and DIPEA in DCM:DMF 1:1 in a fritted SPE tube, for 20 minutes 2 times at room temperature.

**Cleavage**

Peptides were cleaved using TFA:DTT:TIS:$H_2O$ mixture (88:5:2:5) for 3 hours, after which TFA was evaporated and the crude peptide was precipitated in ice-cold diethyl-ether. The precipitate was washed with ether then redissolved in ACN:$H_2O$ mixture and lyophilized.

##### Peptide purification

The following eluents were used for all preparative HPLC purifications and HPLC-UV methods: A: 0.1% TFA/$H_2O$; B: 0.1% TFA/ACN.

$HIF_{786-826}$ and $HIF_{776-826}$ were dissolved in $H_2O$ and purified on a Phenomenex Luna C18 (250 x 10 mm) column using the following gradients: 5 min 0% B, 15 min 20% B, 75 min 50% B, 4 ml/min flow rate; or 5 min 0% B, 15 min 35% B, 75 min 65% B, 4 ml/min flow rate. Purity of the fractions were analysed by HPLC-UV using a Kinetex EVO C18 (5 mm, 100 Å, 250 x 4.6 mm) column and the following gradient: 0 min 5% B, 25 min 80% B, 1 ml/min flow rate.

$CITED_{216}$ and $CITED_{226}$ were dissolved in DMSO and purified on a Phenomenex Luna C4 (250 x 10 mm) column using the following gradient: 0 min 25% B, 5 min 25% B, 15 min 45% B, 70 min 75 %B, 4 ml/min. Fractions containing the peptides were collected, freeze-dried and repurified on the same column or on Kinetex EVO C18 (250 x 10 mm, 100 Å, 5 mm) using the following gradient: 0 min 25% B, 5 min 25% B, 15 min 50% B, 70 min 80 %B, 3 ml/min. Purity of the fractions were analysed using HPLC-UV on an Aeris Widepore C4 (250 x 4.6 mm) column using the following gradient: 0 min 10% B, 25 min 90 % B.

#### Fluorescence anisotropy

##### Direct binding

GSTp300 protein was serially diluted in buffer (25 mM Tris-HCl, 150 mM NaCl, 1mM DTT, pH 7.4) and GFP-HIF-1α (50nM final concentration) was added, the plates were read after 0 min, 60 min, 4 h and 24 h, and incubated at room temperature in between. Each experiment was run in triplicate and the fluorescence anisotropy measured using an EnVision 2103 MultiLabel plate reader (Perkin Elmer) with excitation at 480 nm and emission at 535 nm (30 nm bandwidths). In parallel, a control experiment was performed in which no GFP-HIF was added and the volume made up with additional buffer, this blank was deducted from the raw data for each of the three repeats. The intensity was calculated for each point using eqn (1) and used to calculated anisotropy using eqn (2). From a plot of anisotropy against GST-p300 concentration the minimum and maximum anisotropies were obtained using a logistic sigmoidal fit in OriginPro 8.6. This allowed the conversion to fraction bound (eqn (3)). The data were then fitted using eqn (4) in OriginPro 8.6 to determine the dissociation constant, Kd.

$$I = (2*PG) + S \qquad \text{(eq. 1)}$$

$$r = (S - PG)/I \qquad \text{(eq. 2)}$$

$$L_b = (r - r_{min})/((\lambda(r_{max} - r)) + r - r_{min}) \qquad \text{(eq. 3)}$$

$$y = ((K_d + x + [FL]) - \sqrt{((K_d + x + [FL])^2\ 4x[FL]))/2} \qquad \text{(eq. 4)}$$

r = anisotropy, I = total intensity, P = perpendicular intensity, S = parallel intensity, G = an instrument factor set to 1, $L_b$ = fraction ligand bound, $I = I_{bound}/I_{unbound} = 1$, [FL] = concentration of fluorescent peptide, $K_d$ = dissociation constant, y = $L_b$ multiplied by [FL], x = protein concentration.

#### Isothermal titration calorimetry

Prior to ITC measurement p300 CH1 was dialysed or run on S75 column in the buffer used for titration, and the same buffer was used to dissolve the pure peptides. Experiments with expressed HIF-1α, CITED, CITIF and GFP-HIF-1α variant peptides were performed using a MicroCal iTC200 instrument with 10 µM protein in the cell and 100 µM ligand in the syringe at 37°C, using 2 µl injections and 120 or 180 s spacing between injections. Experiments with the synthetic HIF-1α, CITED and CITIF peptides were carried out using a Microcal VP ITC instrument using 3-5 µM protein in the cell and 40-60 µM ligand in the syringe at 35°C, using 5 µl injections and 180 s spacing between injections, for 40 injections. Titrations were performed in 40 mM Na-phosphate, pH 7.5 100 mM NaCl, 1 mM DTT buffer except for GFP-HIF-1α and variant for which 25 mM Tris-HCl, 150 mM NaCl, 1mM DTT, pH 7.4 buffer was used. Data was analysed using NITPIC[3] and fitted using SEDPHAT[4] to a single binding site model, figures were prepared using GUSSI[5].

##### Competition ITC measurements

Competition ITC measurements were performed in Microcal VP ITC. For competition ITC measurements the P300/ligand complexes were prepared by titrating the appropriate ligand first to p300 in the cell using 3-5 µM protein concentration, the titration stopped when it reached close to saturation conditions (i.e. no significant difference in injection heats). This resulted in 1.15-1.5 excess of ligand compared to p300 in the cell, the recalculated concentrations for each experiment are listed in Table S4. The additional volume was removed from the cell and the complex was titrated with the competitor ligand using 20-50 µM concentration in the syringe, 5 or 10 uL injections with 180-240 s spacing. Background titrations were performed without p300 in the cell and subtracted from the measurements.

**Table S4** Concentrations used in competition ITC experiments.

| | *Cell* | *Syringe* | *Injection volume / spacing* |
|---|---|---|---|
| CITED2$_{224\text{-}259}$ titrated to HIF-1α$_{786\text{-}826}$/P300 complex | 4.5 μM P300<br>5.5 μM HIF$_{786\text{-}826}$ | 50 μM CITED$_{224\text{-}259}$ | 8 μl / 240 s |
| HIF-1α $_{786\text{-}826}$ titrated to CITED2$_{224\text{-}259}$ /P300 complex | 4.4 μM P300<br>5.9 μM CITED$_{224\text{-}259}$ | 50 μM HIF$_{786\text{-}826}$ | 10 μl / 240 s |
| CITED2$_{216\text{-}269}$ titrated to HIF-1α $_{776\text{-}826}$/P300 complex | 2.1 μM P300<br>3.6 μM HIF$_{776\text{-}826}$ | 20 μM CITED$_{216\text{-}269}$ | 10 μl / 240 s |
| HIF-1α $_{776\text{-}826}$ titrated to CITED2$_{216\text{-}259}$ /P300 complex | 5 μM P300<br>5.8 μM CITED$_{216\text{-}269}$ | 50 μM HIF$_{776\text{-}826}$ | 5 μl / 180 s |
| CITED2$_{216\text{-}248}$ titrated to HIF-1α $_{776\text{-}826}$/P300 complex | 4.4 μM P300<br>5.9 μM HIF$_{776\text{-}826}$ | 50 μM CITED$_{216\text{-}248}$ | 10 μl / 240 s |

Integration was performed using NITPIC. In order to fit apparent $K_{\mathrm{D,app}}$ and $\Delta H_{app}$ values data were fitted to a binary interaction model in SEDPHAT. For estimation of $K_{\mathrm{D,app}}$ and $\Delta H_{app}$ based on the individual ligand binding parameters, the following equations were used:

$$K_{app} = K_A \frac{1 + \alpha K_B[B]}{1 + K_B[B]}$$

$$\Delta G_{app} = \Delta G_A - RTln\frac{1 + \alpha K_B[B]}{1 + K_B[B]}$$

$$\Delta H_{app} = \Delta H_A - \Delta H_B \frac{K_B[B]}{1 + K_B[B]} + (\Delta H_B + \Delta\Delta H)\frac{\alpha K_B[B]}{1 + \alpha K_B[B]}$$

where Kapp is the fitted apparent association constant, $K_A$ is the association constant for the ligand in the syringe, $K_B$ is the association constant for the competitor ligand in complex with P300 in the cell, [B] is the concentration of the competitor ligand and α is the cooperativity constant.

Global analysis[6,7] of the competition experiments were performed using different models implemented in SEDPHAT, the best model for each dataset was selected based on the global $\chi^2$ of the fit. For datasets including CITED$_{224\text{-}259}$ interactions the global analysis was performed using the following datasets: HIF$_{786\text{-}826}$ titrated to p300 (B to A); CITED$_{224\text{-}259}$ titrated to p300 (C to A); CITED$_{224\text{-}259}$ titrated to p300/HIF$_{786\text{-}826}$ complex (C to AB) and HIF$_{786\text{-}826}$ titrated to p300/CITED$_{224\text{-}259}$ (B to AC) complex and $K_A$, $\Delta H_A$, $K_B$, $\Delta H_B$ were fitted using the A+B+C <-> AB + C <-> AC + B competition model in SEDPHAT. For datasets including CITED$_{216\text{-}269}$ the global analysis was performed with the following measurements: HIF$_{776\text{-}826}$ titrated to p300 (B to A); CITED$_{216\text{-}269}$ titrated to p300 (C to A); CITED$_{216\text{-}269}$ titrated to p300/HIF$_{776\text{-}826}$ complex (C to AB) and $K_A$, $\Delta H_A$, $K_B$, $\Delta H_B$, cooperativity constant ($\log K_{[AB]C}/K_{AC}$) and $\Delta\Delta H$ were fitted using A+B+C <-> AB + C <-> AC + B <-> ABC triple complex model. Incompetent fraction for p300 was fitted locally for each experiment. 68 % confidence intervals for the fitted values were determined using the automatic confidence interval search implemented in SEDPHAT.

**Co-crystallization**

p300 was mixed with CITIF and the complex was purified by size-exclusion chromatography (S75 26/600 pg column in 20 mM Tris, 150 mM NaCl, 1mM DTT pH 7.5 buffer). Fractions containing the complex were concentrated to a final protein concentration of 5-6 mg/ml. Sparse matrix screening using the JCSG Core suites (Qiagen) was performed with the sitting-drop vapor-diffusion method at 20 °C. Protein was mixed with crystallization solution at a 1:1 ratio with final drop volume of 0.2 µl using NT8® crystallization robot and the Rock Maker® platform (Formulatrix). Initial hit conditions were further optimized by screening HEPES pH 6 – 7.5 in 0.5 pH steps and PEG 6K concentration from 15-40% in 5% steps. Crystals were flash cooled in liquid nitrogen and sent to Diamond Light Source (DLS) for data collection. Data were collected at 100 K. Data were processed with the xia2[8] bundle using DIALS[9] for integration and Pointless[10], Aimless[11] for scaling and merging. Phasing was performed by molecular replacement using Phaser[12] and using 7LVS.pdb as the search model. Initial refinement was done using REFMAC[13] with model building in COOT[14] using the CCP4i2[15] software package. Further refinements and TLS refinement were done in PHENIX[16] and the structures were analysed by Molprobity[17].

**Table S5** Oligonucleotide sequences used for site-directed mutagenesis

| | |
|---|---|
| L795A | 5'-acttcacaatcataactggtcgcctgtggtaatccactttcatc-3' |
| | 5'-gatgaaagtggattaccacaggcgaccagttatgattgtgaagt-3' |
| D799A | 5'-ggagcattaacttcacaagcataactggtcagctgtg-3' |
| | 5'-cacagctgaccagttatgcttgtgaagttaatgctcc-3' |
| E801A | 5'-gctgaccagttatgattgtgcagttaatgctcctatacaag-3' |
| | 5'-cttgtataggagcattaactgcacaatcataactggtcagc-3' |
| N803A | 5'-ccttgtataggagcagcaacttcacaatcataactggtcagctgtg-3' |
| | 5'-cacagctgaccagttatgattgtgaagttgctgctcctatacaagg-3' |
| L818A | 5'-tgatccaaagctctgagtgcttcttcaccctgcagtaggt-3' |
| | 5'-acctactgcagggtgaagaagcactcagagctttggatca-3' |
| L822A | 5'-gagctagttaacttgatccgcagctctgagtaattcttcaccc-3' |
| | 5'-gggtgaagaattactcagagctgcggatcaagttaactagctc-3' |
| D823A | 5'-ctcgagctagttaacttgagccaaagctctgagtaattc-3' |
| | 5'-gaattactcagagctttggctcaagttaactagctcgag-3' |
| V825A | 5'-ggtgctcgagctagttagcttgatccaaagctctg-3' |
| | 5'-cagagctttggatcaagctaactagctcgagcacc-3' |
| L792A | 5'-ctggtcagctgtggtgctccactttcatccattgattgcc-3' |
| | 5'-ggcaatcaatggatgaaagtggagcaccacagctgaccag-3' |
| E817A | 5'-ccaaagctctgagtaatgcttcaccctgcagtagg-3' |
| | 5'-cctactgcagggtgaagcattactcagagctttgg-3' |
| S797A | 5'-gcattaacttcacaatcataagcggtcagctgtggtaatccac-3' |
| | 5'-gtggattaccacagctgaccgcttatgattgtgaagttaatgc-3' |
| Q824A | 5'-gtgctcgagctagttaactgcatccaaagctctgagtaattcttc-3' |
| | 5'-gaagaattactcagagctttggatgcagttaactagctcgagcac-3' |
| L818A, L822A | 5'-gagctagttaacttgatccgcagctctgagtgcttcttcaccctgcagtaggttt-3' |
| | 5'-aaacctactgcagggtgaagaagcactcagagctgcggatcaagttaactagctc-3' |
| L818A L822A V825A | 5'-ggtgctcgagctagttagcttgatccgcagctctgagtgcttcttcaccctgcagta-3' |
| | 5'-tactgcagggtgaagaagcactcagagctgcggatcaagctaactagctcgagcacc-3' |
| L818A L822A D823A | 5'-gaaacctactgcagggtgaagaagcactcagagctgcggctcaagttaactagctcgagc-3' |
| | 5'-gctcgagctagttaacttgagccgcagctctgagtgcttcttcaccctgcagtaggtttc-3' |
| L795A D799A | 5'-tataggagcattaacttcacaagcataactggtcgcctgtggtaatccactttcatcc-3' |
| | 5'-ggatgaaagtggattaccacaggcgaccagttatgcttgtgaagttaatgctcctata-3' |
| L795A D799A E801A | 5'-ccttgtataggagcattaactgcacaagcataactggtcgcctgtggtaatccactttcatc-3' |
| | 5'-gatgaaagtggattaccacaggcgaccagttatgcttgtgcagttaatgctcctatacaagg-3' |
| L795A D799A E801A N803A | 5'-tctgctgccttgtataggagcagcaactgcacaagcataactggtcgcctgtggtaatccactttcatcc-3' |
| | 5'-ggatgaaagtggattaccacaggcgaccagttatgcttgtgcagttgctgctcctatacaaggcagcaga-3' |

### Peptide characterization data

#### **HIF-1α$_{786\text{-}826}$** (Synthetic)

Ac-SMDESGLPQLTSYDCEVNAPIQGSRNLLQGEELLRALDQVN-$NH_2$

Em: 4543.185, MW: 4546.001

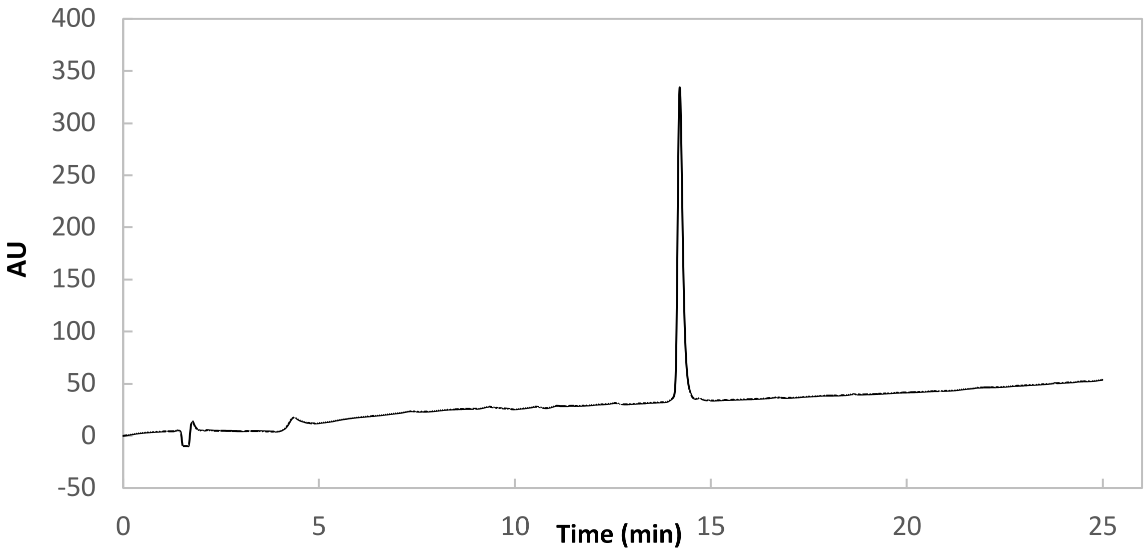

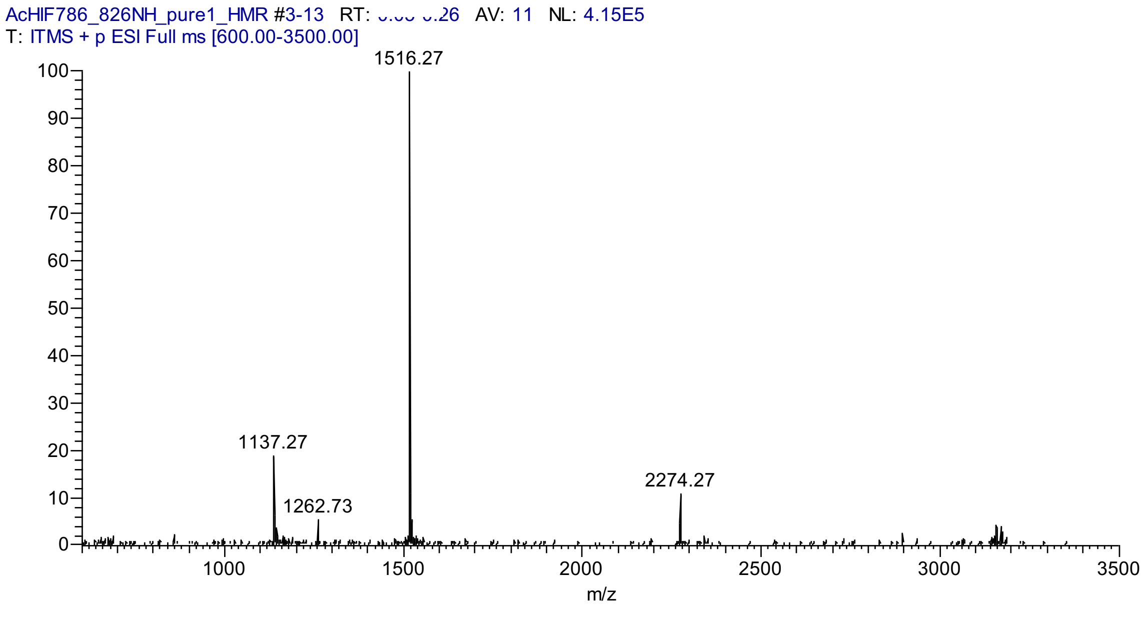

**HIF-1α$_{776-826}$** (Synthetic)

Ac-SDLACRLLGQSMDESGLPQLTSYDCEVNAPIQGSRNLLQGEELLRALDQVN-$NH_2$

Em: 5599.724;

Mw: 5603.236

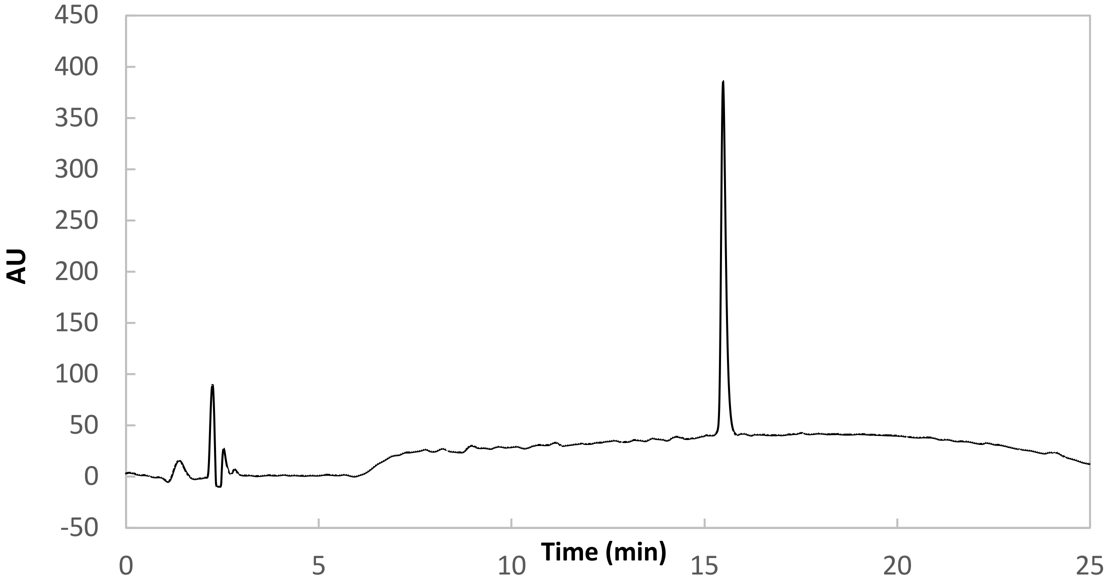

98s

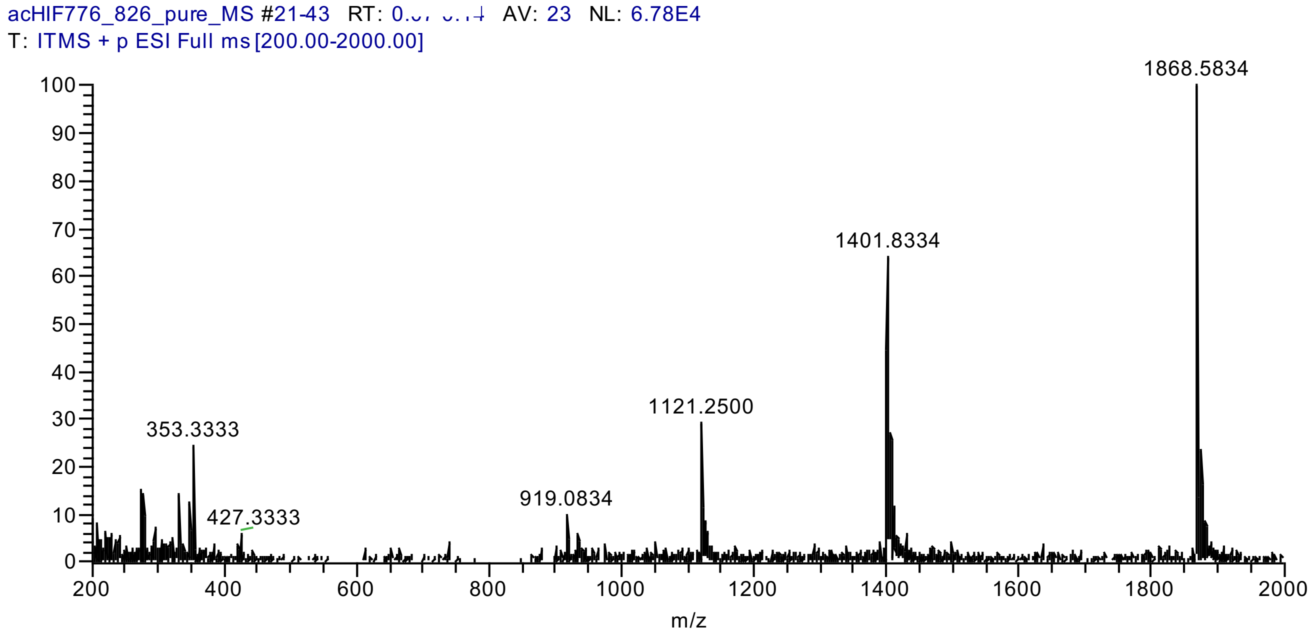

**CITED2$_{224\text{-}259}$** (Synthetic)

Ac-DEEVLMSLVIEMGLDRIKELPELWLGQNEFDFMTDF-$NH_2$

Em: 4342.078

Mw: 4344.963

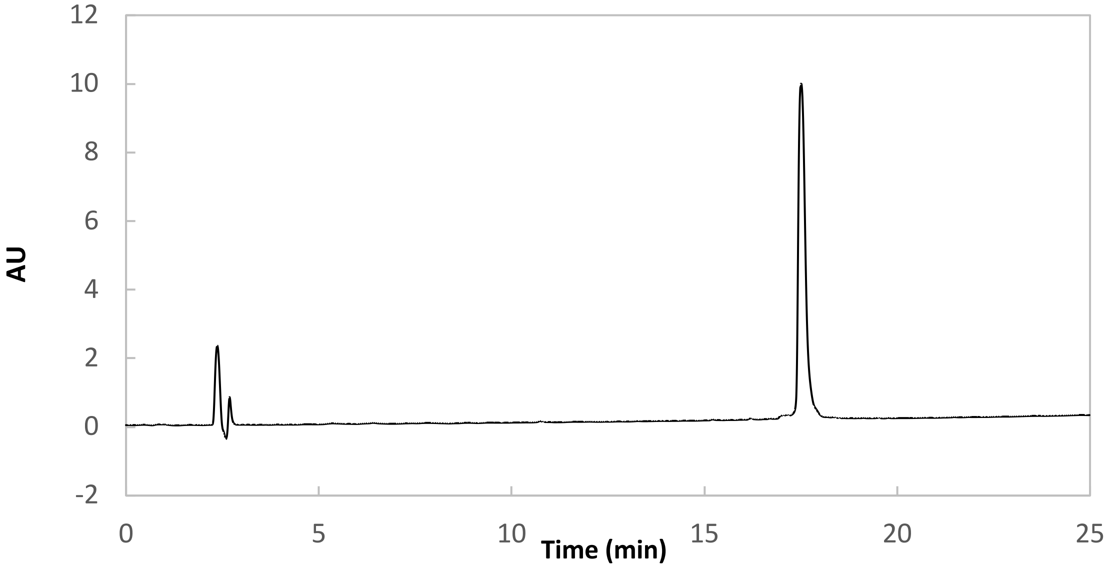

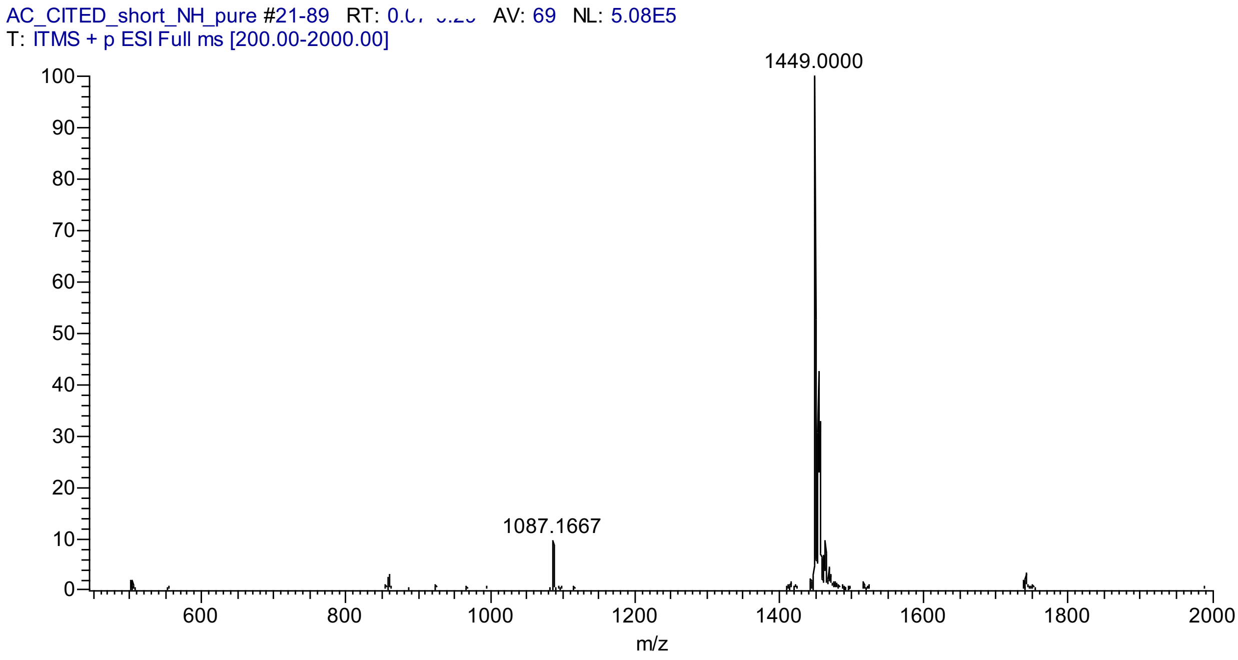

#### CITED2$_{216\text{-}269}$ (Synthetic)

Ac-NVIDTDFIDEEVLMSLVIEMGLDRIKELPELWLGQNEFDFMTDFVCKQQPSRVS-$NH_2$

Em: 6372.104

MW: 6376.276

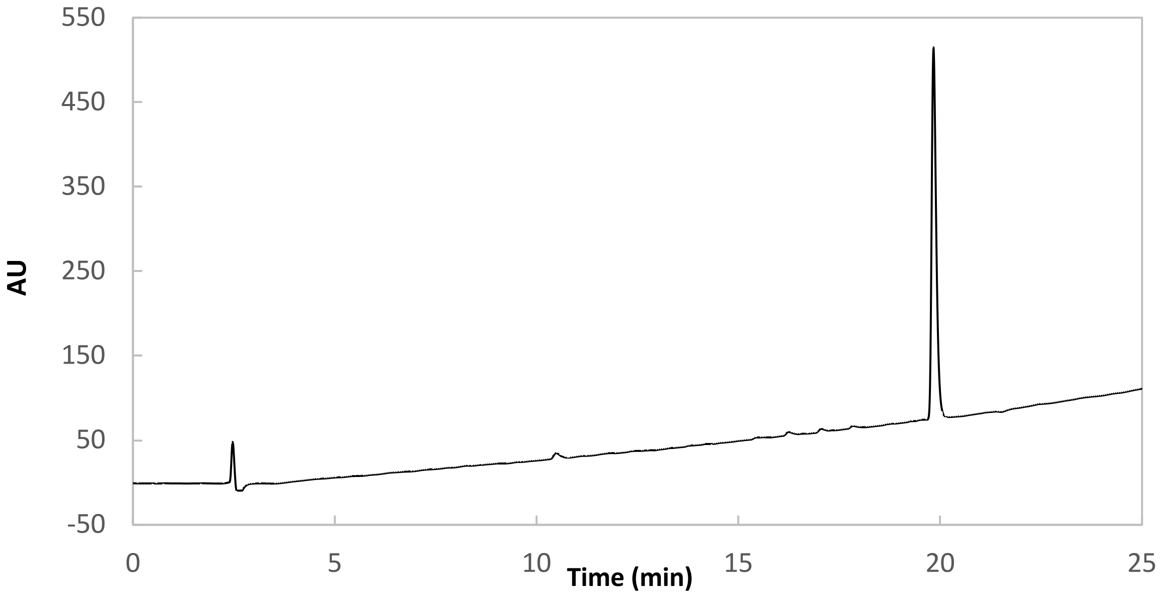

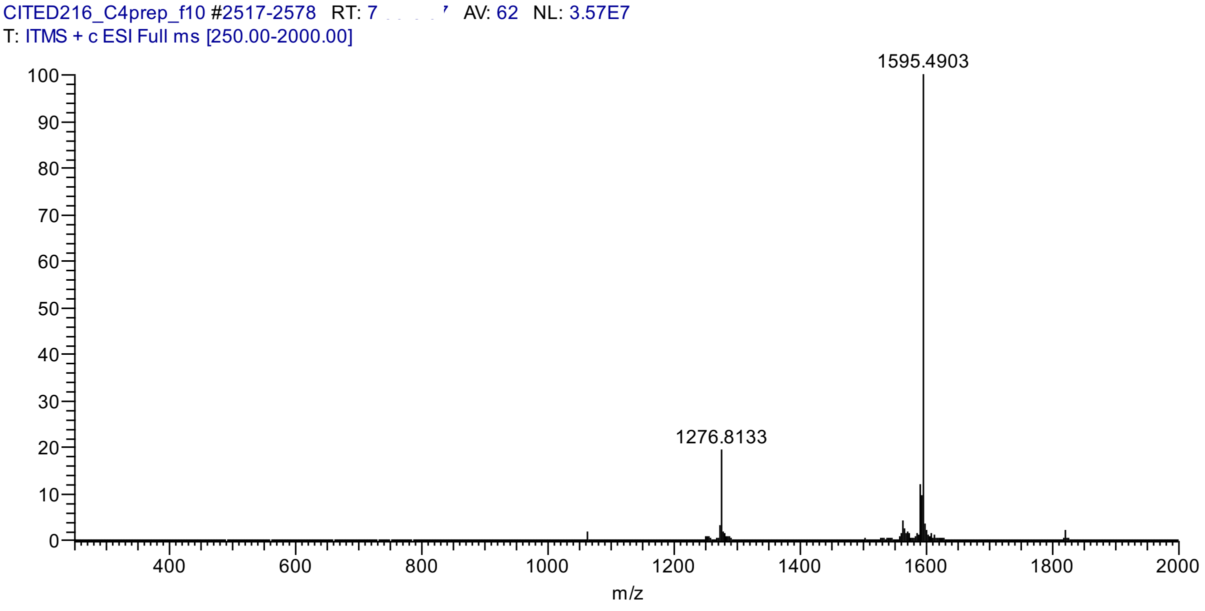

#### CITIF (synthetic)

Ac-DEEVLMSLVIEMGLDRIKELPQLTSYDCEVNAPIQGSRNLLQGEELLRALDQVN-$NH_2$

Em: 6137.102

Mw: 6140.975

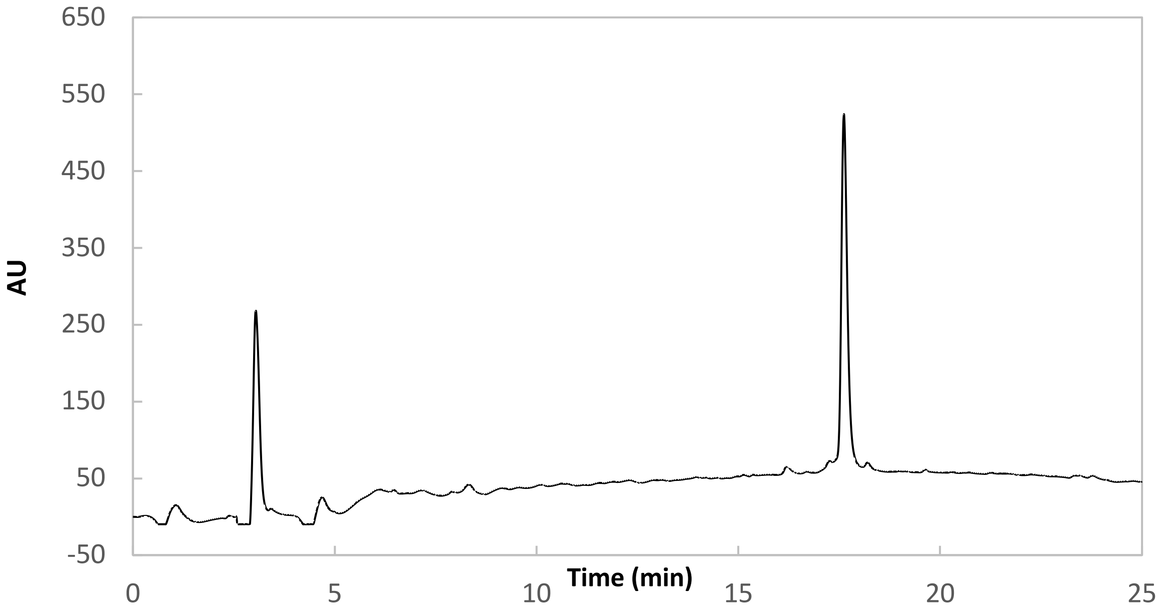

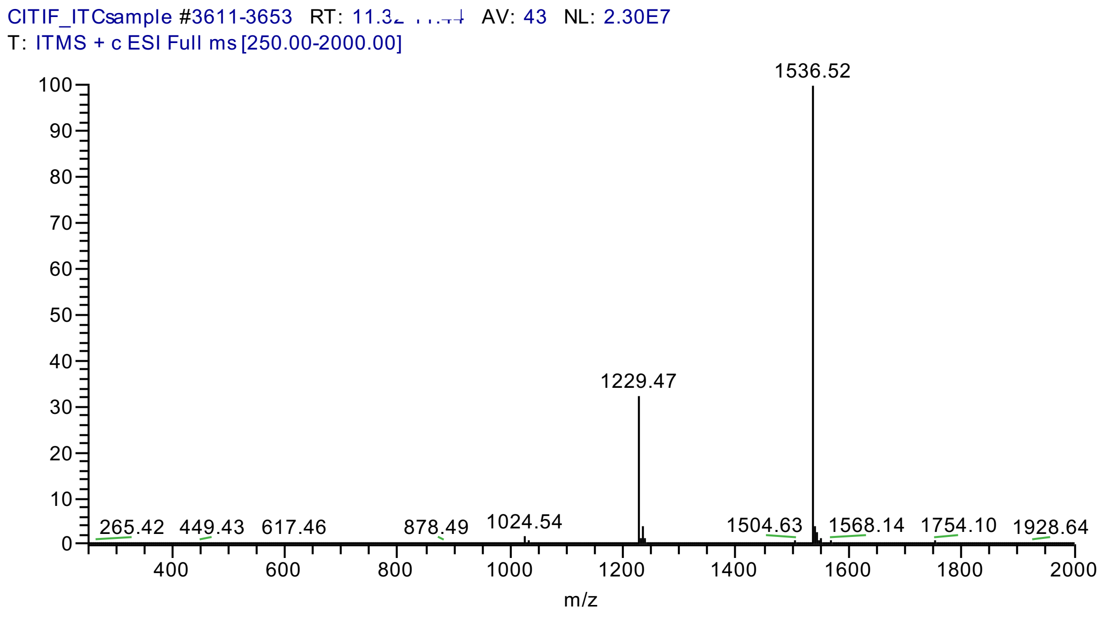

**CITED2$_{216\text{-}248}$**

Ac-NVIDTDFIDEEVLMSLVIEMGLDRIKELPELW-$NH_2$

Em: 3814,93

Mw: 3817,41

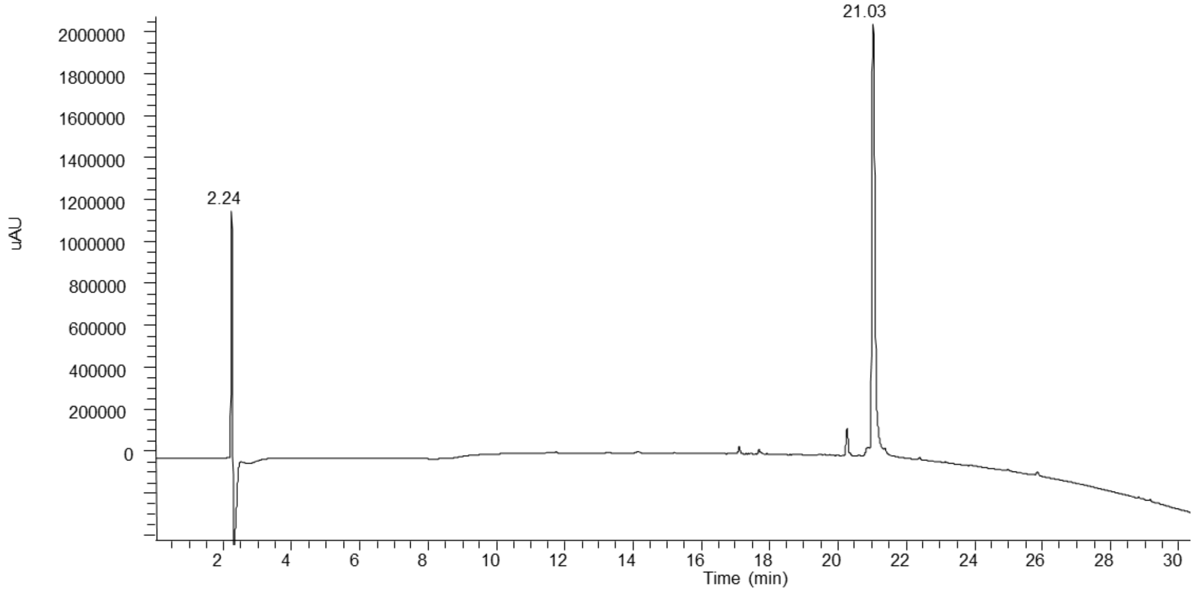

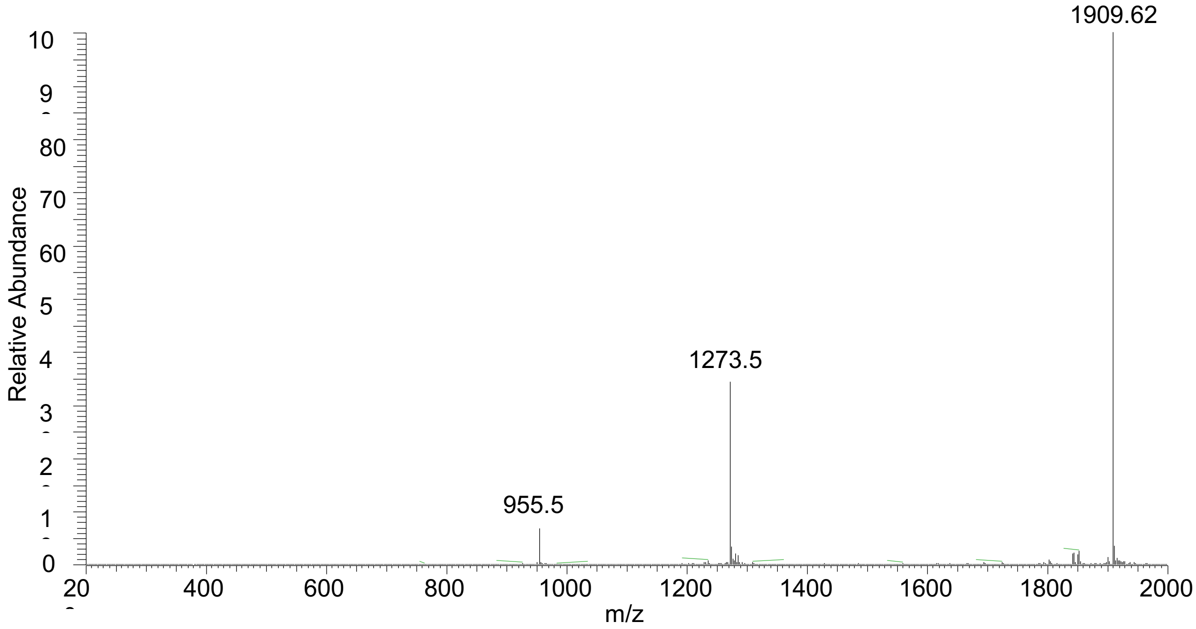

**CITED2$_{224\text{-}248}$**

Ac-DEEVLMSLVIEMGLDRIKELPELW-$NH_2$

Em: 2897,48

Mw: 2899,40

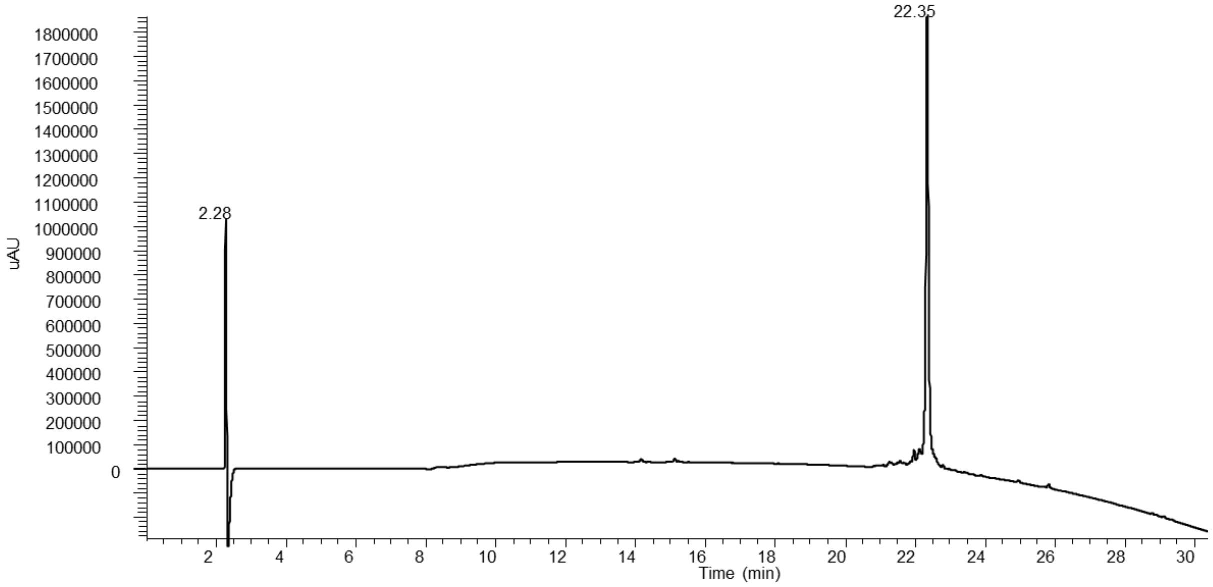

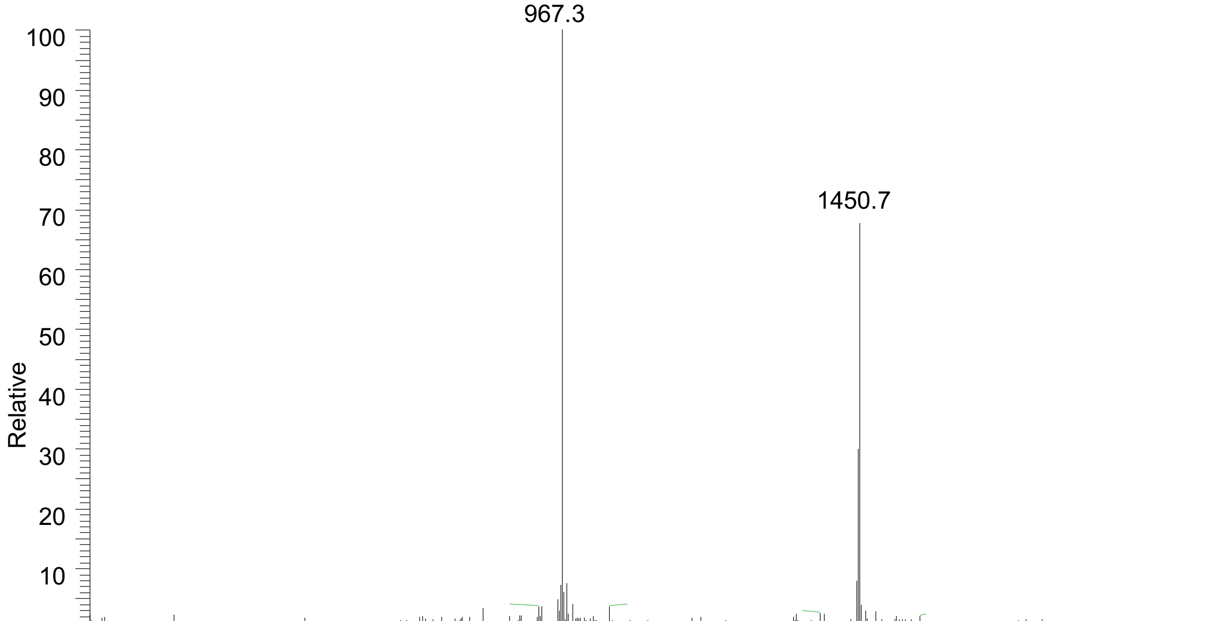

### Characterization data for the expressed HIF-1α and CITED and CITIF constructs

#### HIF-1$\alpha_{776-826}$

GPGSSDLACRLLGQSMDESGLPQLTSYDCEVNAPIQGSRNLLQGEELLRALDQVN-OH

Em: 5856.825

MW: 5860.481

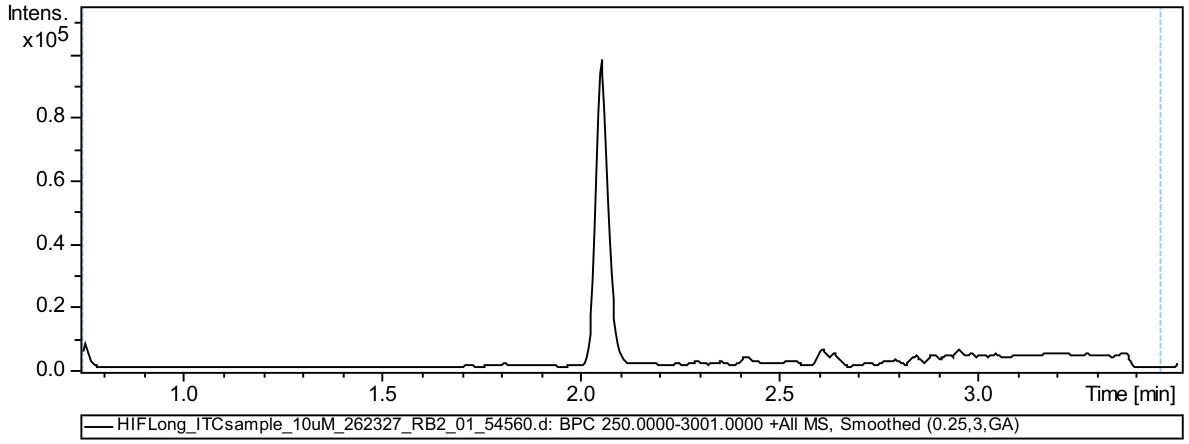

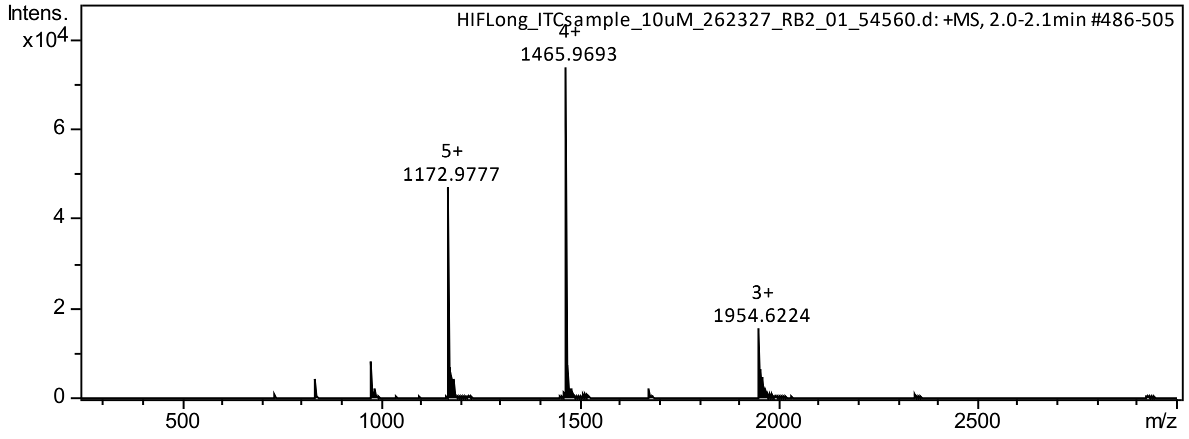

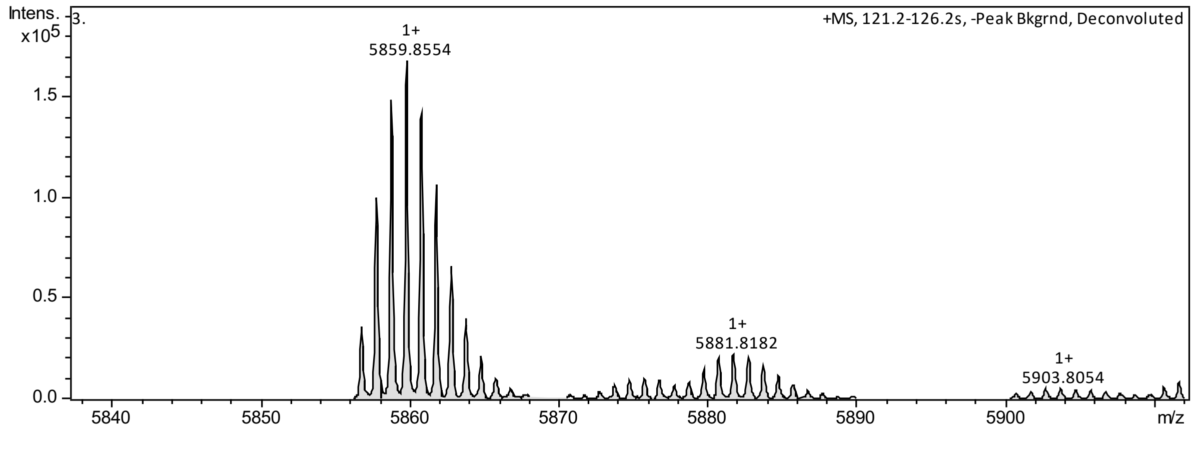

**CITED2**$_{\mathbf{224\text{-}259}}$ (expressed)

GPGSDEEVLMSLVIEMGLDRIKELPELWLGQNEFDFMTDF-OH

MW: 4602.208

Em: 4599.179

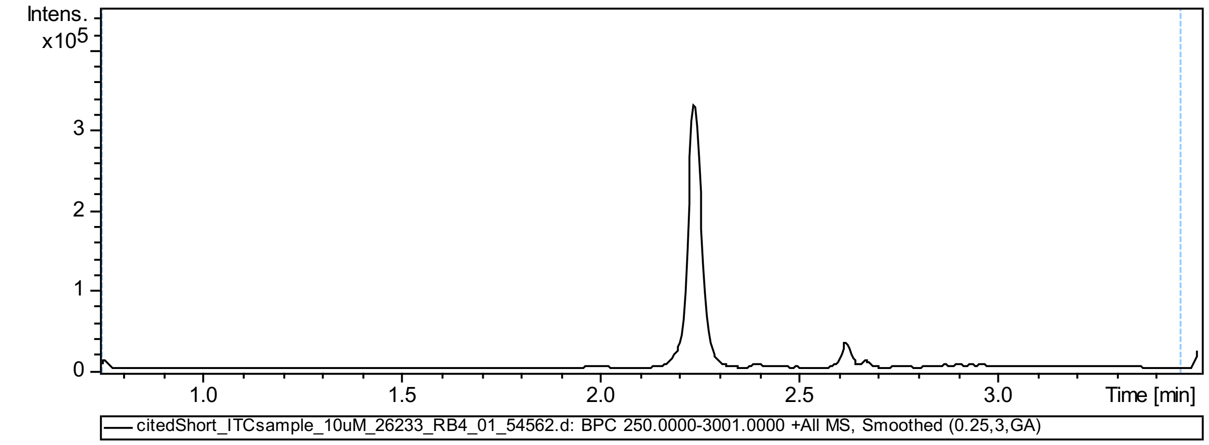

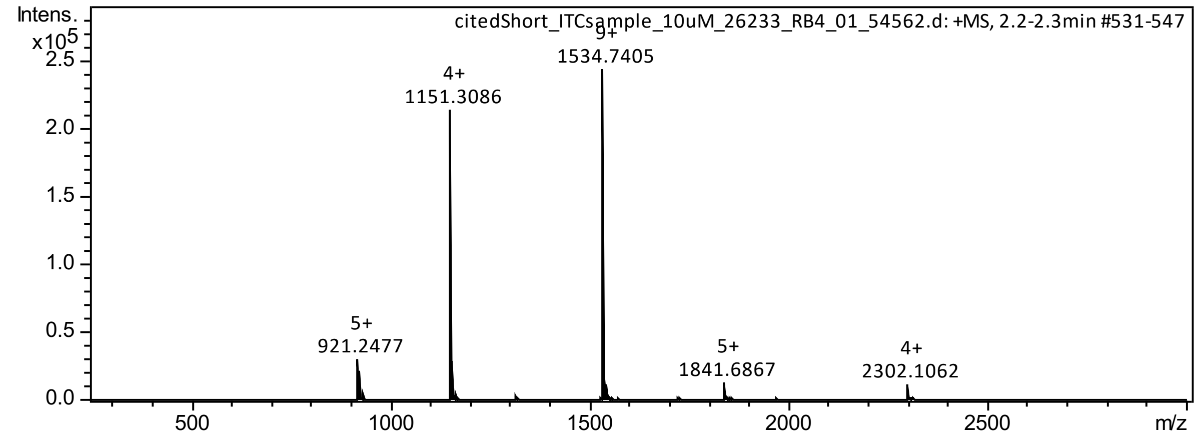

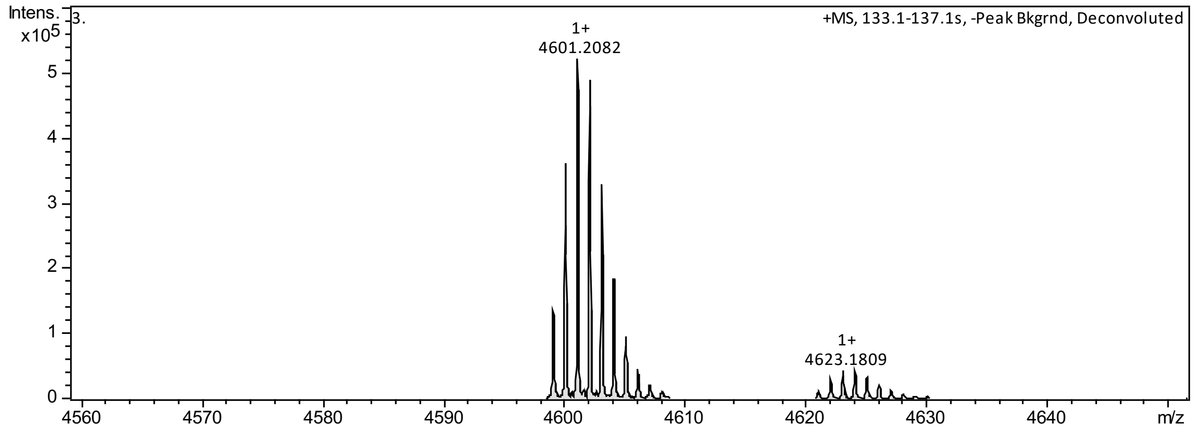

**CITED2$_{216\text{-}269}$** (expressed)

GPGSNVIDTDFIDEEVLMSLVIEMGLDRIKELPELWLGQNEFDFMTDFVCKQQPSRVS-OH

MW: 6633.521

Em:6629.205

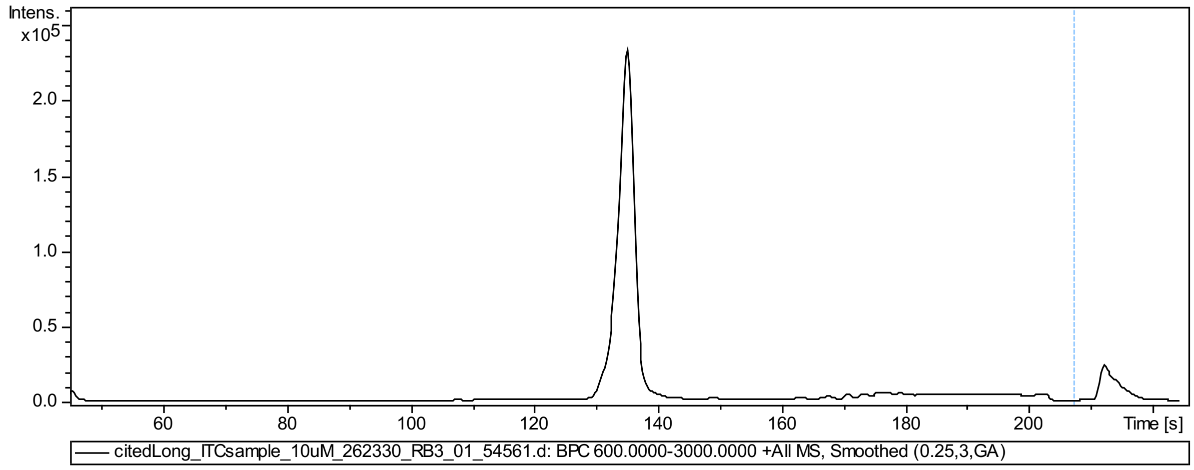

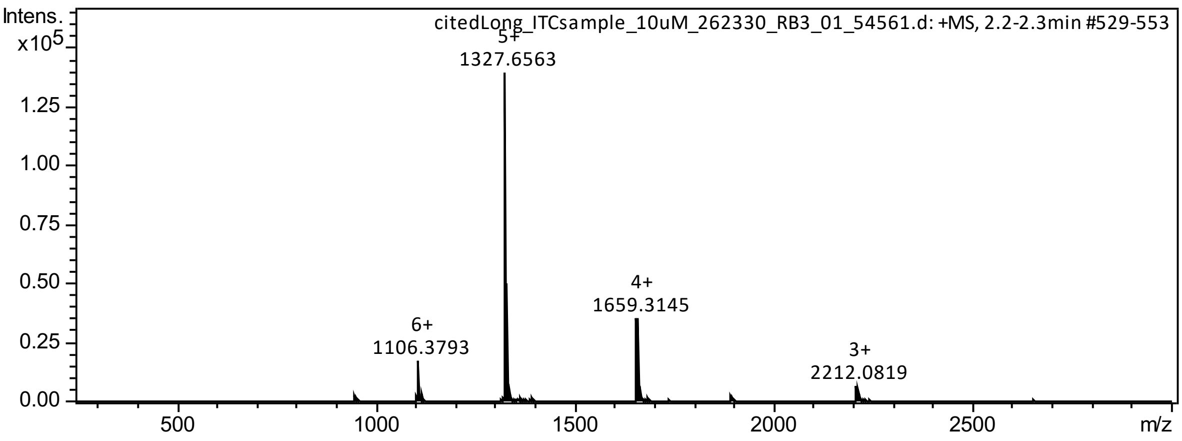

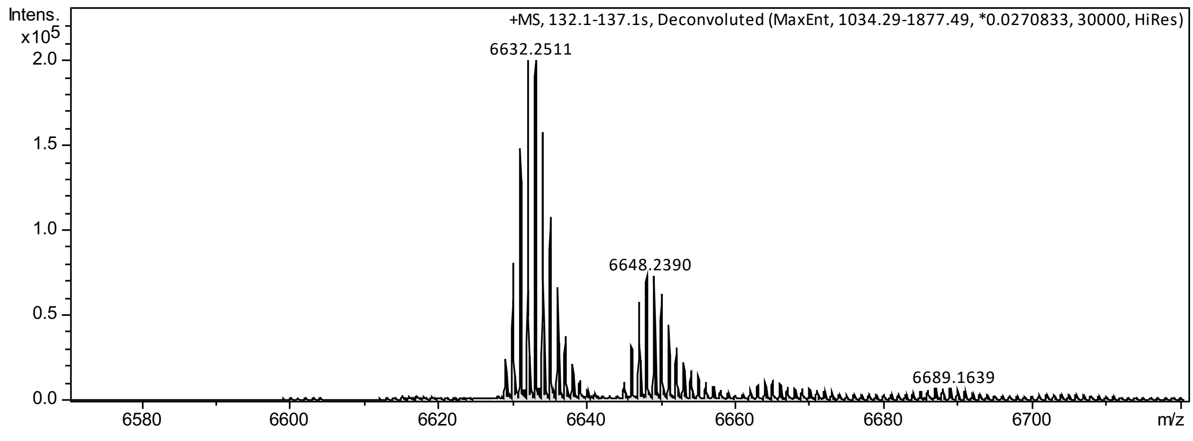

**CITIF (expressed)**

GPGSDEEVLMSLVIEMGLDRIKELPQLTSYDCEVNAPIQGSRNLLQGEELLRALDQVN-OH

MW: 6398.220

Em: 6394.203

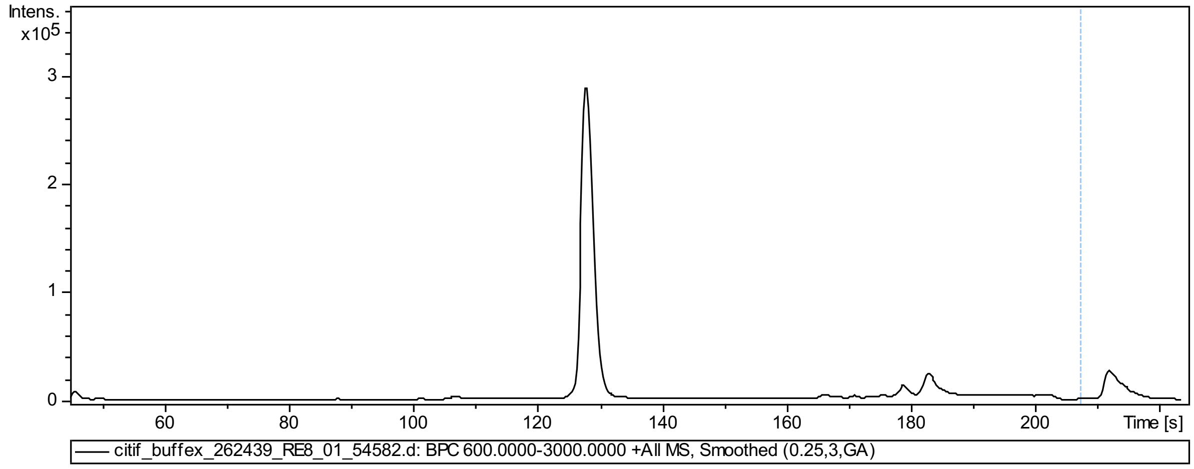

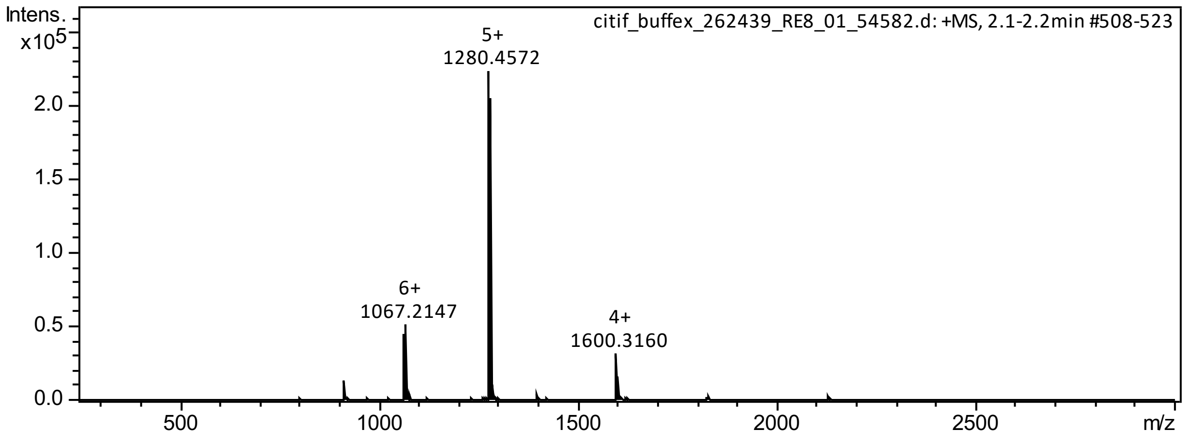

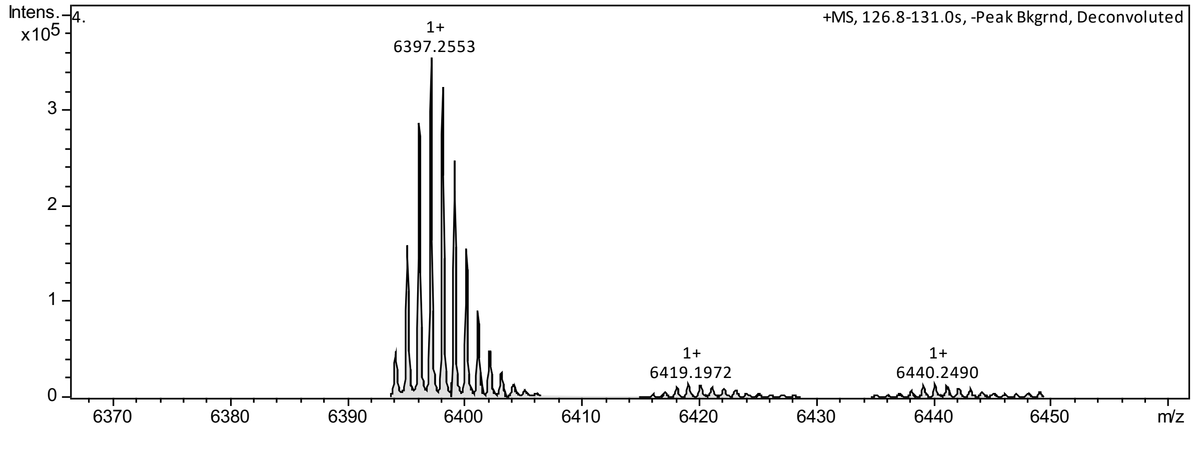
